# Arabidopsis GALACTURONOSYLTRANSFERASE (GAUT) 1 synthesizes a homogalacturonan tightly bound to the cell wall and required for cell expansion

**DOI:** 10.64898/2025.12.22.695998

**Authors:** Melani A. Atmodjo, Robert A. Amos, Kalina T. Haas, Alexis Peaucelle, John Glushka, Li Tan, Ian M. Black, P. Jarod Glatz, Darryn C. Amanda, Ioana Emerich, Joshua Indech, Sivakumar Pattathil, Sindhu Kandemkavil, Stefan Eberhard, Parastoo Azadi, Michael G. Hahn, Debra Mohnen

**Author notes:** To whom correspondence should be addressed: Debra Mohnen, Department of Biochemistry and Molecular Biology, Complex Carbohydrate Research Center, The University of Georgia, 315 Riverbend Rd. Athens, GA 30602-4712. Mascoma LLC (Lallemand Inc.), 67 Etna Rd., Lebanon, NH 03766, USA. The author responsible for distribution of materials integral to the findings presented in this article in accordance with the policy described in the Instructions for Authors (https://academic.oup.com/plcell/pages/General-Instructions) is Debra Mohnen.

## Abstract

Arabidopsis GALACTURONOSYLTRANSFERASE1 (GAUT1) synthesizes homogalacturonan (HG), the most abundant pectin in growing plant cells. GAUT1 has the greatest *in vitro* enzyme activity of the six confirmed Arabidopsis HG biosynthetic GAUTs, but its biological activity remains elusive. Here we show that Arabidopsis *GAUT1* homozygous mutants have a severe dwarfed seedling phenotype, survive several weeks as 2 to 3 mm seedlings, and have severely reduced shoot and root growth and hypocotyl epidermal, cortex and endodermal cell size. *gaut1-1* pollen tubes are shorter than WT with increased bursting. Complementation of homozygous *gaut1-1* with *GAUT1* coding sequence driven by the *GAUT1* promoter restored WT-like growth. The extreme dwarf phenotype of homozygous *gaut1-1* seedlings precluded their use for detailed cell wall analysis, thus suspensions cultures were produced from callus generated from mutant and WT seedlings. Homozygous *gaut1-1* suspension cells were smaller than WT with ∼30% reduced wall GalA content compared to WT. Sequential extraction of the walls with increasingly harsh solvents and sugar composition analysis revealed reduced GalA content in only the 4M KOH post-chlorite fraction, indicating that GAUT1-synthesized HG was held tightly in the wall by direct or indirect hydrogen bonding and/or oxidation-sensitive linkages. Treatment of wall fractions with endopolygalacturonase to hydrolyze HG and gel electrophoretic separation of hydrolysates exposed an HG-associated doublet band markedly downregulated in the homozygous *gaut1-1* 4M KOH post-chlorite fraction and to a lesser extent in 4M KOH and sodium chlorite fractions. NMR analysis identified the band as rhamnogalacturonan (RG)-II. Super resolution microscopy using anti-HG antibodies showed that, compared to WT, the homozygous *gaut1-1* hypocotyl epidermal and callus cells had reduced content and length of HG nanofilaments, HG fibers associated with cell expansion in Arabidopsis. The results demonstrate that GAUT1-synthesized HG resides in a tightly-cell-wall-bound, RG-II-containing polymer required for HG nanofilament formation and seedling cell expansion.

## INTRODUCTION

The plant cell wall pectic polysaccharides are ancient and essential glycopolymers present throughout the plant kingdom including in extant charophytic green algae (Domozych et al., 2014), indicating functions in plants for over 400 million years. In plants, pectins function in cell expansion, cell:cell adhesion, fruit development, pollen tube growth and plant defense, to name a few (reviewed in Delmer et al. (2024)). Pectins are cell wall polysaccharides that contain 4-linked D-galacturonic acid (GalA), a glyco signature in the two pectic backbones: homogalacturonan (HG) [-α-1,4-D-GalA-α-1,4-D-GalA-]_n_ and rhamnogalacturonan (RG) [-α-1,4-D-GalA-α-1,2-L-Rha-]_n_. The pectic backbones may contain methyl and/or acetyl moieties and have glycosyl side branches yielding the pectin glycan domains HG, rhamnogalacturonan I (RG-I) and rhamnogalacturonan II (RG-II). These pectin domains are covalently connected in the wall via their backbones resulting in diverse complex pectic cell wall heteroglycans whose specific structures and functions are not well understood (Mohnen et al., 2024). HG is the most abundant pectin in the walls of growing and dividing cells in non-graminaceous flowering plants. It is also present in plant cells and tissues with secondary walls that predominantly contain cellulose and hemicelluloses. In such walls, HG is primarily located in the middle lamella where it functions in cell:cell adhesion via HG-calcium-HG salt bridges and as the backbone of RG-II which forms RG-II borate-diester pectin crosslinks (Mohnen et al., 2024).

HG is synthesized by α-1,4-<u>ga</u>lact<u>u</u>ronosyltransferases (GAUTs) (Sterling et al., 2006; Atmodjo et al., 2011; Hao et al., 2014; Amos et al., 2018; Voiniciuc et al., 2018; Engle et al., 2022). GAUT1 was discovered by isolation of Arabidopsis (*Arabidopsis thaliana*) suspension culture cell proteins that co-purified with HG:α-1,4-galacturonosyltransferase (HG:GalAT) activity and mass spectrometric sequencing of the partially purified proteins (Sterling et al., 2006). *In vivo*, Arabidopsis GAUT1 is devoid of its transmembrane domain due to post-translational N-terminal cleavage and held in the Golgi apparatus, the site of pectin synthesis, by enzyme complex formation with its homologs GAUT5, GAUT6, or GAUT7 (Atmodjo et al., 2011; Lund et al., 2020). *In vitro,* Arabidopsis GAUT1 has been shown to be the catalytic subunit of the GAUT1:GAUT7 complex, which elongates HG acceptors into high molecular weight polymers and also initiates HG synthesis in the absence of exogenous acceptors using uridine diphosphate (UDP)-GalA as both an acceptor and donor substrate (Amos et al., 2018).

Bioinformatics analysis revealed that Arabidopsis GAUT1 exists in a 15-member GAUT family (Sterling et al., 2006), an unexpected finding since only three GAUTs would be predicted to be required to synthesize HG, the backbone of RG-II, and HG-RG-I heteroglycans (Mohnen et al., 2024). Currently, six Arabidopsis GAUTs have been biochemically confirmed to have HG:GalAT activity (GAUTs 1, 4, 10, 11, 13, 14) (Sterling et al., 2006; Amos et al., 2018; Biswal et al., 2018b; Voiniciuc et al., 2018; Engle et al., 2022). As mentioned above, three additional GAUTs (GAUTs 5, 6, 7) function to anchor GAUT1 in the Golgi via formation of protein complexes (Atmodjo et al., 2011; Lund et al., 2020). The existence of so many catalytic GAUTs leads to the hypothesis that different GAUTs synthesize HG species in unique cell wall polymers.

One way to test this hypothesis is to analyze GAUT mutants and identify the polymer/polymers affected. Cell wall analyses of T-DNA insertion mutants of 13 out of the 15 Arabidopsis *GAUT* gene family members have been reported (Caffall et al., 2009). Distinct patterns of altered sugar composition compared to wild-type (WT) were identified in cell walls of mutants of eight different *GAUT* genes (i.e., *GAUTs 6*, *8*, *9*, *10*, *11*, *12*, *13* and *14*), suggesting a unique function for each GAUT. T-DNA insertion mutants were not available at that time for *GAUT1* and *GAUT4,* suggesting lethal homozygous phenotypes. As this work was being written for publication, a report of a GAUT1 CRISPR-knockout (KO) suspension culture line with a 2012 bp deletion of the GAUT1 gene (from exon 5 to exon 11) was published with the expressed goal of generating a low aggregation cell suspension culture (Frankevich et al., 2025). The authors reported abnormal homozygous and heterozygous GAUT1 CRISPR-KO seedlings that did not form normal stem and leaves, confirming the importance of the *GAUT1* gene for plant growth and development. Suspension cell cultures generated from the GAUT1 CRISPR-KO homozygous seedlings were found to be darker in color with reduced growth rates, larger cell aggregates and cells of smaller size compared to the vector control. The authors reported that the pectin content in the GAUT1 CRISPR-KO cells was equivalent to the vector control based on a commercial Pectin Identification Assay Kit, a finding that is at odds with the results we report below in this paper.

Here we took a molecular genetic approach to determine the biological and structural function of Arabidopsis GAUT1. To this end, we isolated and characterized Arabidopsis *gaut1* T-DNA insertion mutants. The *gaut1* homozygous seeds had reduced germination frequency and those that germinated grew as extremely dwarfed tiny seedlings due to defective cell expansion. The stunted seedlings yielded no mature plants, and the tiny dwarf phenotype did not provide the amount of material required for cell wall biochemical analyses. Thus, homozygous *gaut1-1/gaut1-1* mutant suspension cultured cells were used as a whole plant proxy, revealing a ∼30% reduction in cell wall GalA content in comparison to WT. Unexpectedly, GAUT1-synthesized HG was found to reside in a fraction tightly bound into the cell wall that required harsh chemical extraction for release from the wall. Evidence was obtained indicating that the HG synthesized by GAUT1 resides in a specific HG:RG-II pectic polymer that is downregulated in the *gaut1-1/gaut1-1* homozygous mutant compared to WT. Super resolution microscopy using anti-HG monoclonal antibodies revealed that *gaut1-1/gaut1-1* homozygous cell walls have reduced HG nanofilament content in comparison to WT cell walls. Taken together, the results reported here support the hypothesis that GAUT1 synthesizes a particular HG that is held tightly in the wall, is associated with specific RG-II, is required for pectin nanofilament formation, and that without this HG cells cannot expand/elongate properly.

## RESULTS

### *gaut1-1/gaut1-1* knock-out mutant seedlings have a stunted, growth-arrested phenotype

As of the writing of this manuscript, five germplasm lines were listed for Arabidopsis *GAUT1* (*At3g61130*) on the TAIR website (Table S1). The SAIL_1162_A03 line harbors a single T-DNA insertion in the 5’-untranslated region (UTR) of the *GAUT1* gene, but genotyping identified only WT individuals. The WiscDsLoxHs190_06B line also carries a single T-DNA insertion in the 5’-UTR of the *GAUT1* gene; genotyping of this line revealed homozygous individuals but these grew indistinguishably from WT, suggesting that the T-DNA insertion did not affect *GAUT1* expression. The GABI-KAT line 470H07-019028, as obtained, has one additional T-DNA insertion elsewhere in genome, which we removed by backcrossing to WT and screening of progeny over four generations. From this, a line harboring a single T-DNA insertion within *GAUT1* exon 10 was isolated and designated as the *gaut1-1* mutant line (Figure 1A). The SALK_063880 and SALK_130652 lines each carried two additional T-DNA insertions elsewhere in the genome. Seedlings homozygous for the T-DNA insertion within the *GAUT1* gene in these two lines were identified and found to have stunted growth characteristics comparable to those of *gaut1-1* (see below). These two alleles were designated the *gaut1-2* and *gaut1-3* mutant lines, respectively (Figure 1A). Since comparable phenotypes were obtained in three different T-DNA insertion mutants, we focused on characterization of the *gaut1-1* mutant and its complemented line to probe the function of GAUT1-synthesized HG in Arabidopsis.

**Figure 1.**
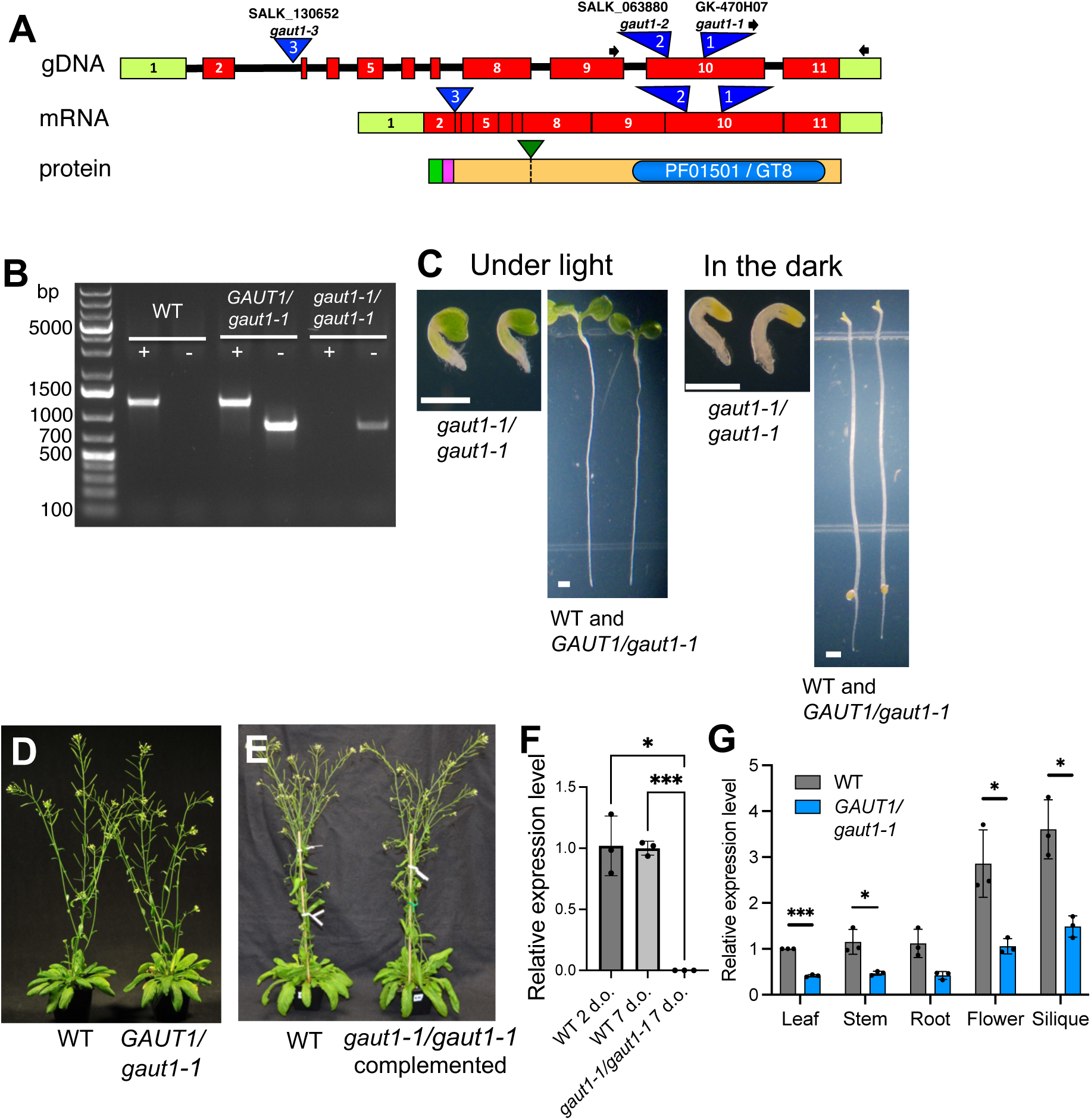
Arabidopsis *gaut1-1* mutant with a single T-DNA insertion within the *GAUT1* gene was isolated from GABI-KAT line 470H07-019028. (A) Gene, transcript, and protein structures of Arabidopsis *GAUT1* (*At3g61130*). For gene and transcript: red boxes denote exons; light green boxes denote 5’- and 3’-untranslated regions (UTRs); black lines denote introns; blue down-facing triangles indicate the location of the T-DNA insertion within the exon 10 of *GAUT1* for *gaut1-1* and *gaut1-2*, and within the intron 2 for *gaut1-3*; black arrows indicate the position and direction of gene- and T-DNA-specific PCR primers used to perform genotyping of *gaut1-1*. For the GAUT1 protein: green box denotes the predicted *N*-terminal cytosolic domain; magenta box denotes the predicted trans-membrane domain; blue oval denotes the GT8 domain; green down-facing triangle denotes the *in vivo* cleavage site of Arabidopsis GAUT1 protein reported previously (Atmodjo et al., 2011). (B) PCR-based genotyping of WT, heterozygous, and homozygous *gaut1-1*, performed on 7-day-old seedlings. PCR reactions using primer sets targeting WT and mutant loci are in lanes indicated with (+) and (-) signs, respectively. (C) Homozygous *gaut1-1/gaut1-1* was recovered as severely stunted seedlings compared to the normal-looking seedlings of WT and heterozygous *GAUT1/gaut1-1*. Seven-day-old seedlings grown under light or in the dark are shown. Scale bars = 1 mm. (D) Mature 8-week-old WT and *GAUT1/gaut1-1* plants show no visible difference in growth. (E) Mature 8-week-old WT and complemented *gaut1-1/gaut1-1* plants show comparable growth. (F and G) Relative transcript levels of *GAUT1* as determined by quantitative RT-PCR of RNA extracted from (F) whole seedlings of WT (2 and 7 days old) and *gaut1-1/gaut1-1* (7 days old), and (G) various tissues from 8-week-old WT and *GAUT1/gaut1-1* plants. The expression of *GAUT1* in WT seedlings (in F) and leaves (in G) is set to 1, and *Actin2* was used as the reference gene. Data are means ± standard deviation (n=3). Asterisks indicate significant difference as analyzed by one-way ANOVA (in F) or Student’s t-test (in G) (* *P* < 0.05, *** *P* < 0.001).

The relatively high level and widespread transcript expression of Arabidopsis *GAUT1* (Atmodjo et al., 2011) hinted at an essential function for normal plant growth and development and suggested that its absence would result in lethality or observable phenotypes. When segregating seeds from *GAUT1/gaut1-1* heterozygous plants were germinated on solid medium plates, as early as 3 days after germination, homozygous mutant (*gaut1-1/gaut1-1*) individuals could be identified as tiny, very stunted seedlings that grew no larger than a few millimeters, in contrast to the normal-looking WT and heterozygous mutant seedlings (Figure 1B, C). No mature plants were recovered from the stunted *gaut1-1/gaut1-1* seedlings, while heterozygous *GAUT1/gaut1-1* grew into mature plants indistinguishable from WT (Figure 1D, Figure S1A-D). Re-introduction of the *GAUT1* coding sequence driven by the *GAUT1* promoter (*ProGAUT1*) into the *GAUT1/gaut1-1* background followed by genotyping of the T1 generation plants resulted in the recovery of mature complemented *gaut1-1/gaut1-1* homozygous mutant plants that grew comparable to WT (Figure 1E, Figure S1E-G), confirming that the observed mutant phenotypes were caused by the *GAUT1* mutation. Quantitative PCR analysis of RNA from 7-day-old *gaut1-1/gaut1-1* homozygous seedlings and from mature *GAUT1/gaut1-1* heterozygous plants revealed a complete absence of *GAUT1* transcript in the *gaut1-1/gaut1-1* homozygous seedlings, and >50% reduction in transcript level in *GAUT1/gaut1-1* heterozygous plants, compared to WT (Figure 1F, G; Figure S2). Interestingly, there seemed to be a feedback response to the mutation in *gaut1-1/gaut1-1* seedlings based on the significant increases observed in transcript levels of multiple *GAUT1*-related gene family members (i.e., *GAUTs 3, 4, 6, 13*, and *15*; Figure S2) compared to WT.

### *gaut1-1/gaut1-1* seedling stunted growth is due to defective cell expansion and elongation

The basis of the dwarfed phenotype of the *gaut1-1/gaut1-1* mutant was explored via microscopic examination of organ and cell size in the shoot and root tissues. Both shoot and root growth were compromised in *gaut1-1/gaut1-1* stunted seedlings compared to WT, as evident from the significantly shorter hypocotyl and root length in both light- and dark-grown seedlings (Figure 2A-D). Using scanning electron micrographs of seedlings grown under light (Figure 2E-H), we determined the numbers and size of the epidermal cells lining the entire length of the hypocotyl from top to bottom of WT-like (genotype WT or heterozygous *GAUT1/gaut1-1*) and homozygous *gaut1-1/gaut1-1* seedlings. The cell numbers stayed relatively the same from day 3 through day 10 after germination and were comparable between WT-like and *gaut1-1/gaut1-1* seedlings (Figure 2I), consistent with the reported insignificant or absence of cell division during the Arabidopsis hypocotyl growth phase (Gendreau et al., 1997). In contrast, the length and width of the cells in *gaut1-1/gaut1-1* seedlings were significantly reduced by on average 55.0 (± 6.1) and 21.8 (± 9.4) %, respectively (Figure 2J, K), compared to WT, suggesting defective cell elongation and expansion. The *gaut1-1/gaut1-1* seedling hypocotyl epidermal cells were also noticeably misshapen and not as neatly organized compared to WT, and some cells developed root hairs ectopically (Figure 2H). Epidermal cells of the roots also had compromised cell elongation in the *gaut1-1/gaut1-1* mutant, as indicated by the compacted distribution of root hairs in *gaut1-1/gaut1-1* seedlings. This is consistent with *GAUT1* transcript expression along the length of roots being highest at the elongation zone (Figure S3), imparting a critical function in root cell elongation.

**Figure 2.**
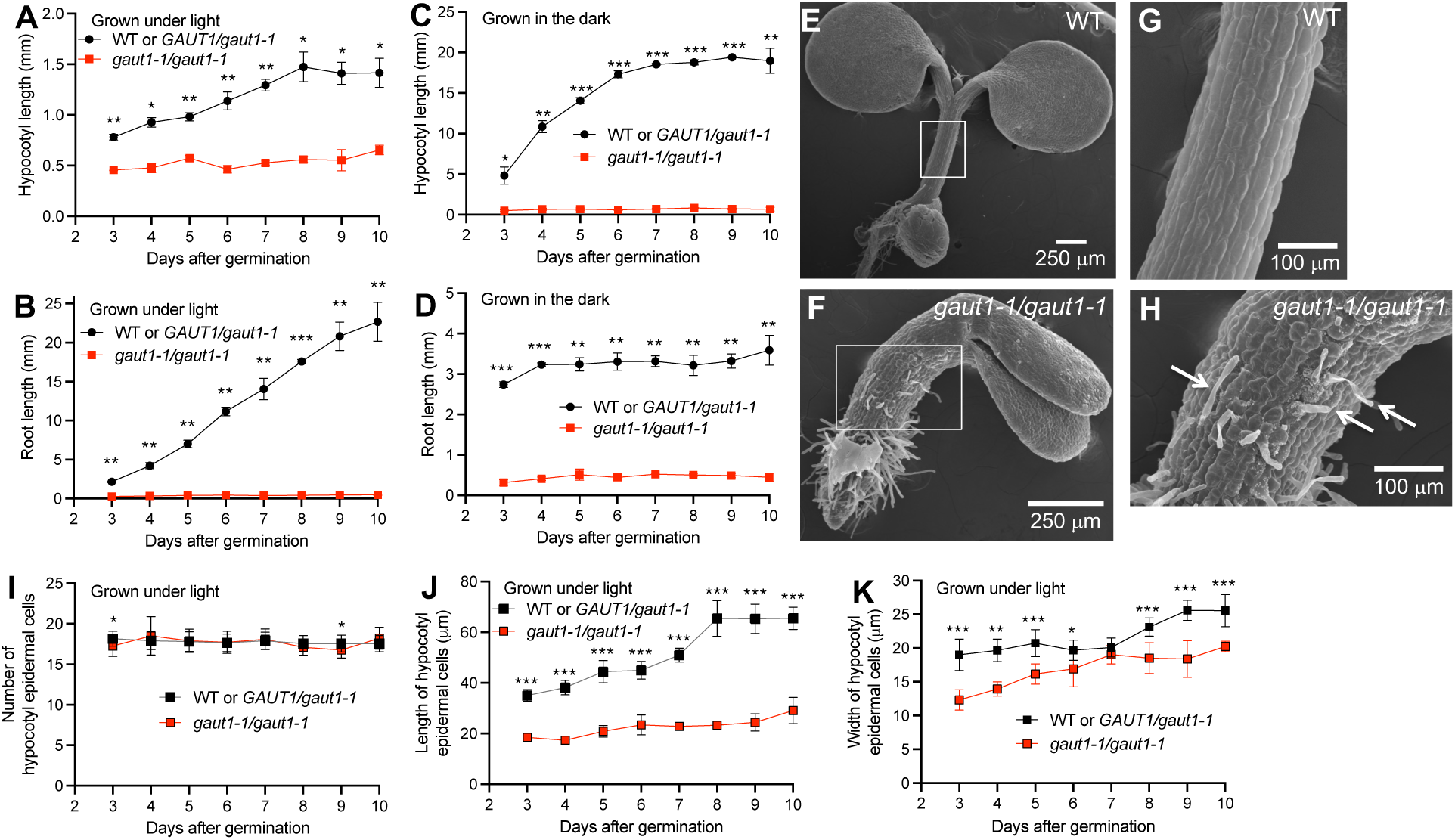
Hypocotyls and roots of *gaut1-1/gaut1-1* homozygous seedlings are significantly shorter than those of WT and *GAUT1/gaut1-1* heterozygous due to smaller cell size. (A-B) Length of hypocotyl (A) and root (B) in seedlings grown under light. (C-D) Length of hypocotyl (C) and root (D) in seedlings grown in the dark. Data are means ± standard deviation from two independent experiments with n ≥ 21 for WT or *GAUT1/gaut1-1* seedlings, and n = 2 to 12 for *gaut1-/-* seedlings in each experiment. Asterisks indicate significant difference as analyzed by Student’s t-test (\**P* < 0.05, \*\**P* < 0.01, *** *P* < 0.001). (E and F) Scanning electron micrographs of 7-day-old seedlings of WT and *gaut1-1/gaut1-1*, respectively, grown under light. (G) and (H) are close-up views of the area indicated by white boxes in (E) and (F), respectively. Arrows in (H) indicate ectopic root hairs growing out of *gaut1-1/gaut1-1* seedling hypocotyl. (I) Numbers, (J) average length, and (K) average width of individual epidermal cells in a single cell file along the length of elongating hypocotyl of seedlings grown under light. Measurement was carried out on two adjacent cell files for (I) and (J), and on one single cell file for (K), of 10 WT or *GAUT1/gaut1-1*, and 4-8 *gaut1-1/gaut1-1* seedlings (except for Day 4 after germination for which only 2 *gaut1-1/gaut1-1* seedlings were analyzed). Data are means ± standard deviation from two independent experiments. Asterisks indicate significant difference as analyzed by Student’s t-test (\**P* < 0.05, \*\**P* < 0.01, *** *P* < 0.001).

We then inspected other aspects of seedling growth that might be affected by the *GAUT1* mutation. The organizational arrangement of the epidermal, cortex, endodermal, and vascular cells in cross sections of seedling hypocotyls appeared comparable between *gaut1-1/gaut1-1* and WT (Figure 3A), although measurement of the cell size from this transversal point of view again showed markedly smaller cell sizes in *gaut1-1/gaut1-1* seedlings compared to WT (Figure 3B), which is consistent with the defective cell expansion described above. We then tested for effects on cell:cell adhesion by immersing etiolated seedlings in ruthenium red solution. Whereas the positive control mutant seedlings of *QUASIMODO1 (QUA1/GAUT8)*, whose hypocotyl epidermal cell:cell adhesion is known to be defective (Bouton et al., 2002), had hypocotyls stained red throughout, those of both WT and *gaut1-1/gaut1-1* stunted seedlings remained white indicating normal cell:cell adhesion (Figure 3C). Finally, when kept on medium plates for an extended period (2-3 weeks), the *gaut1-1/gaut1-1* seedlings appeared to develop multiple protuberances from the apical meristem, which appeared like tiny leaves unable to expand to the proper size and whose epidermal cells looked swollen compared to the WT (Figure 3D-F). This observation suggests that, despite the absence of *GAUT1*, cell division and differentiation are relatively unaffected in the *gaut1-1/gaut1-1* mutant, while developing organs cannot grow to full size due to compromised cell expansion and elongation. Taken together, the above results strongly support a critical function of *GAUT1* in cell expansion and elongation, but not in tissue organization, cell:cell adhesion, cell division, and cell differentiation.

**Figure 3.**
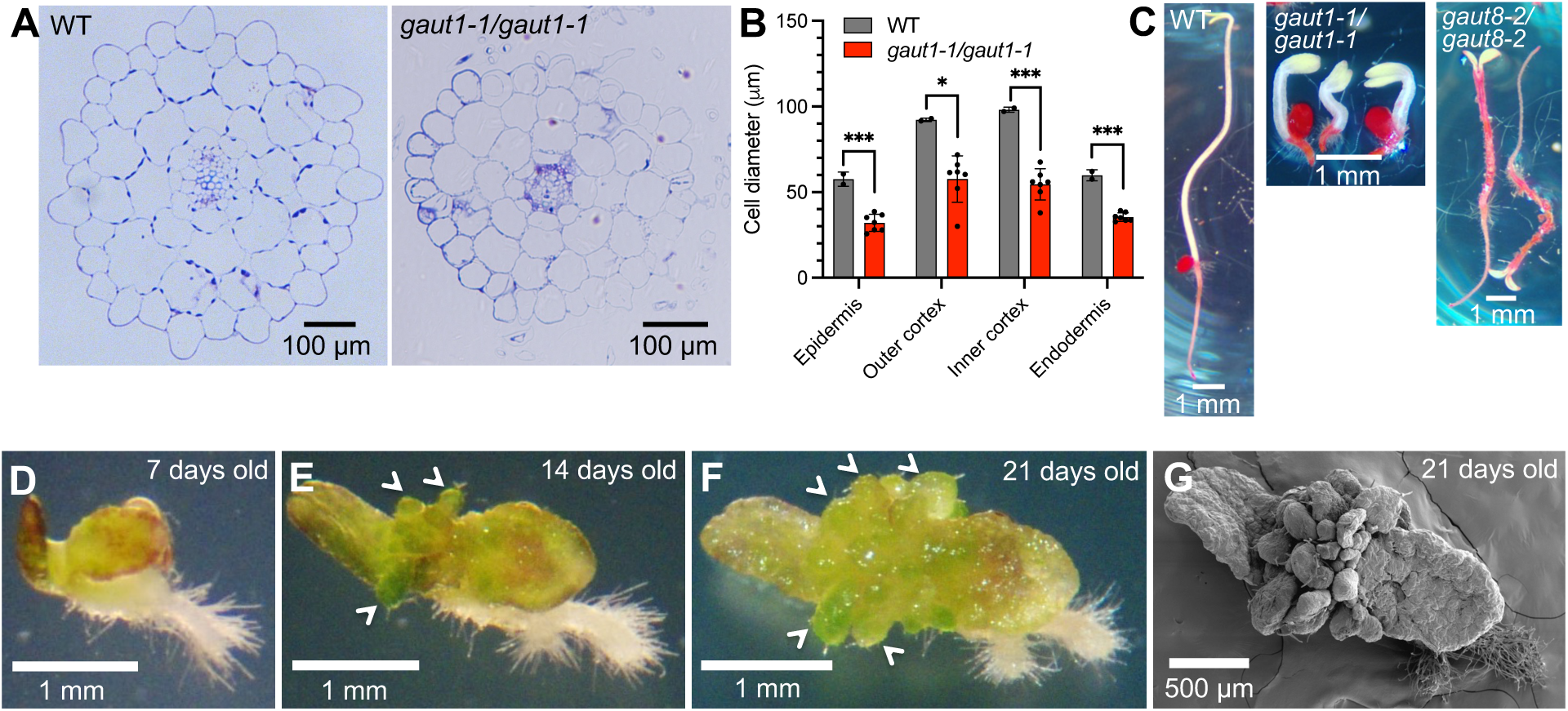
The *gaut1-1* homozygous mutation does not affect tissue organization, cell:cell adhesion, cell division, or cell differentiation, but does affect cell size. (A) Toluidine-blue stained hypocotyl cross sections of 7-day-old WT and *gaut1-1/gaut1-1* seedlings showed comparable tissue organization. (B) Diameter of epidermal, cortex, and endodermal cells shown in (A). Data are means ± standard deviation from two WT and seven *gaut1-1/gaut1-1* seedlings. At least 26 epidermal, 11 outer cortex, 8 inner cortex, and 8 endodermal cells were measured for each seedling. Asterisks indicate significant difference as analyzed by Student’s t-test (* *P* < 0.05, *** *P* < 0.001) (C) Ruthenium red staining of 7-day-old WT and *gaut1-1/gaut1-1* seedlings showed no dye penetration into the hypocotyls, in contrast to the positive control *gaut8-2/gaut8-2* (SALK_039214) mutant whose seedling hypocotyls stained red due to cell:cell adhesion defect. Shown are representatives of three independent experiment (with ≥ 14 mutant seedlings tested in each) with similar results. (D-F) Dissecting scope analysis of *gaut1-1/gaut1-1* seedling development over a 3-week period at (D) 7, (E) 14, and (F) 21 days after germination. White arrow heads indicate stunted leaves. (G) Scanning electron micrograph of the 21-day-old *gaut1-1/gaut1-1* seedling shown in (F).

### *GAUT1* mutation reduces fertility and perturbs embryo/seed development

The *gaut1-1/gaut1-1* homozygous seedlings were observed at a lower frequency than expected for normal segregation (Table 1, 13.3% (363/2728)). We thus investigated whether GAUT1 was critical for fertility. Analysis of the segregating progenies of self-pollinated *GAUT1/gaut1-1* heterozygous plants showed a WT : *GAUT1/gaut1-1* : *gaut1-1/gaut1-1* ratio of 1:1.5:0.4, significantly skewed from the expected 1:2:1 ratio (Table 1). Reciprocal crosses between WT and *GAUT1/gaut1-1* heterozygous plants revealed 28.5% reduced female and 56% reduced male gametophytic transmission compared to the WT (Table 1). These results indicate a function of GAUT1 in reproductive development and fertility.

**Table 1.** Genetic segregation analysis of Arabidopsis *gaut1-1* mutant. All observed ratios are significantly different from the expected ones (ξ_2_, *P* < 0.001). Scoring of the normal-looking WT and heterozygous seedlings versus the dwarfed homozygous ones was by visual observation at day 5 after germination on medium without antibiotic. Scoring of WT versus heterozygous seedlings was by moving the 5-day-old normal-looking seedlings onto medium containing antibiotic sulfadiazine and counting susceptible (Sulf^S^) versus resistant (Sulf^R^) individuals at day 14. Abbreviations: WT – wild type; HT – heterozygous; HM – homozygous. TE_f_ and TE_m_ – female and male transmission efficiency, respectively, which is calculated as (Sulf^R^/Sulf^S^) x 100%. † - combined WT and heterozygous individuals; ‡ - homozygous counts include both germinating seedlings (shown by the number in brackets) and non-germinating seeds.

|  | Ratio |  |  |  |  |  |
| --- | --- | --- | --- | --- | --- | --- |
|  | Expected |  |  | Observed |  |  |
| Selfed *GAUT1/gaut1-1* | WT | HT | HM | W T | HT | HM‡ |
| • analysis of bulked seeds | 3† |  | 1 | 2728† |  | 427 (363) |
| • analysis of seeds from individual siliques | 1 | 2 | 1 | 522 | 790 | 230 (115) |
|  | Expected ratio |  |  | Observed |  |  |
| Back-cross to WT | WT |  | HT | WT | HT |  |
| • ♀ *GAUT1/gaut1-1* X ♂ WT | 1 |  | 1 | 1685 | 1205 | TE<sub>f</sub> = 71.5% |
| • ♀ WT X ♂ *GAUT1/gaut1-1* | 1 |  | 1 | 1323 | 582 | TE<sub>m</sub> = 44.0% |

With male gametophytic transmission being more severely impeded, we examined pollen and pollen tubes to determine the underlying cause. Pollen grains from *GAUT1/gaut1-1* heterozygous plants, which would be expected to be at least 50% normal, appeared to form normally with no apparent loss of viability when stained using fluorescein-diacetate compared to those from WT plants (Figure 4A). There was no difference in the pollen germination rate between WT and *GAUT1/gaut1-1*-derived pollen grains, nor in the percentage of pollen that began to germinate but stopped with only very short pollen tubes (i.e. less than pollen length). However, there was a significantly higher proportion of bursting pollen tubes and fewer pollen with tubes that elongated normally in pollen derived from *GAUT1/gaut1-1* compared to WT (Figure 4B). Importantly, the average tube length of the pollen that elongated normally was significantly reduced for pollen from *GAUT1/gaut1-1* plants compared to WT (Figure 4C). Taken together, the defective pollen tube of the *GAUT1/gaut1-1*-derived pollen is consistent with a flaw in polarized cell elongation of the developing pollen tubes.

**Figure 4.**
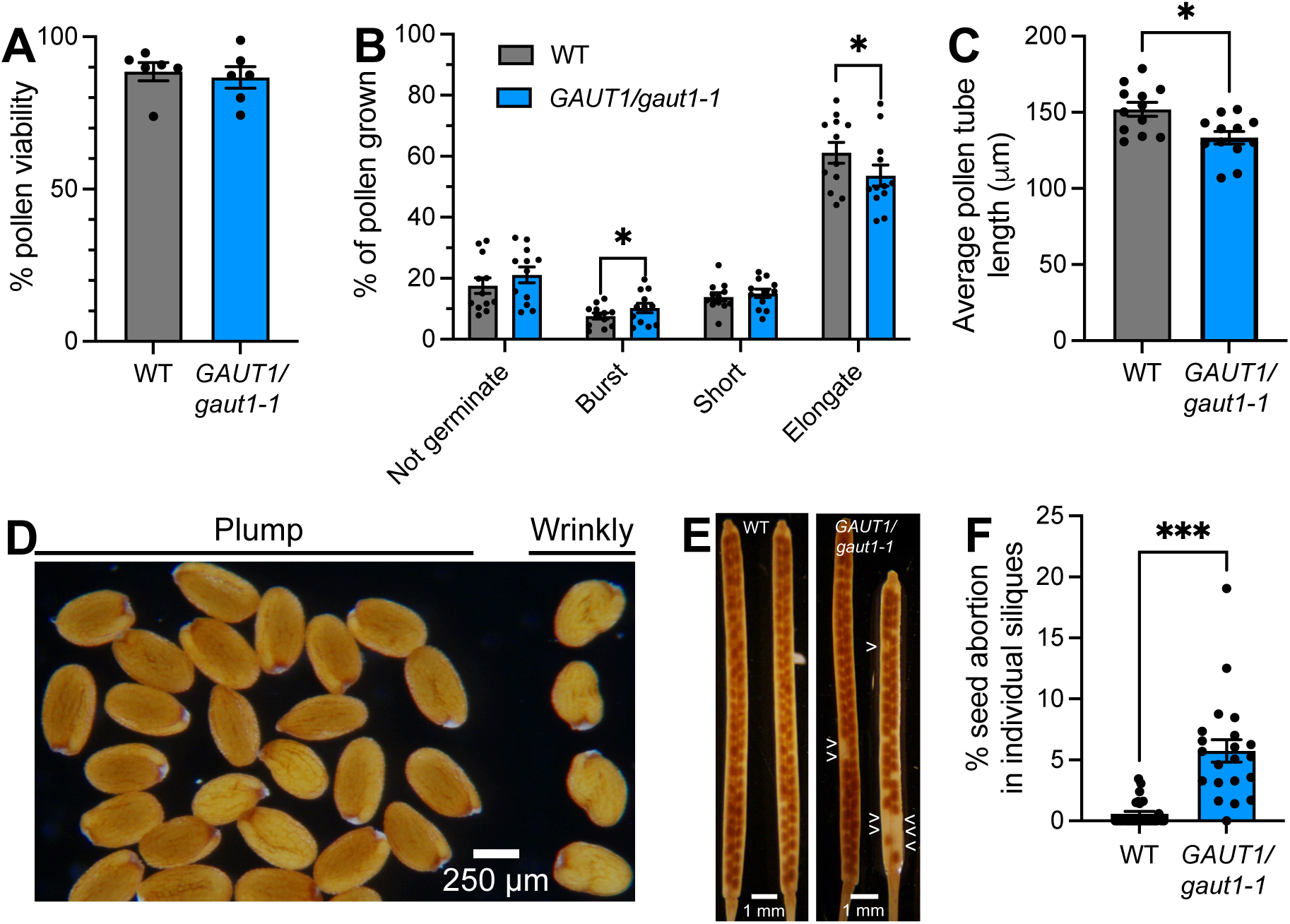
*gaut1-1* homozygous mutation affects reproduction and seed development. (A)Viability of pollen from flowers of WT and *GAUT1/gaut1-1* plants, measured as percentage of pollen grains that fluoresced upon staining with fluorescein diacetate. Data are means ± standard error of six independent observations (n=6), in each of which >400 pollen grains from each genotype were scored. (B) Germination and pollen tube growth performance and (C) pollen tube length of pollen grains from WT and *GAUT1/gaut1-1* flowers, observed after incubation for 4 hours on pollen germination medium *in vitro*. Data are means ± standard error from four technical and three biological replicates (n=12), with the numbers of pollen scored ≥ 300 for (B) and ≥ 100 for (C) in each replicate. Asterisks indicate significant difference as analyzed by Student’s t-test (* *P* < 0.05). (D) A proportion of seeds from *GAUT1/gaut1-1* selfed plants have wrinkly, underdeveloped appearance. (E) Reduced seed set was observed in *GAUT1/gaut1-1* siliques, compared to WT. White arrow heads denote empty spots in the siliques due to seed abortion. (F) Percentage of aborted seeds in WT versus *GAUT1/gaut1-1* siliques. Data are means ± standard error (n ≥ 20 siliques scored for each genotype). Asterisks indicate significant difference as analyzed by Student’s t-test (*** *P* < 0.001).

In addition to the fertility defects, we observed that stunted homozygous seedlings appeared to arise predominantly from underdeveloped seeds with a somewhat shrunken/wrinkly appearance (90.4% (47/52)) rather than from normal plump-looking seeds (7.8% (23/295)) (Figure 4D). Siliques from *GAUT1/gaut1-1* heterozygous plants also had significantly higher incomplete seed sets due to seed abortion compared to WT (Figure 4E and F). These results indicate that the absence of *GAUT1* may cause *gaut1-1/gaut1-1* homozygous individuals to fail during multiple points of seed and seedling development including during vegetative growth of the embryo, seed maturation, seed germination and seedling growth, all of which contribute to the lower-than-normal frequency of the *gaut1-1/gaut1-1* individuals.

### *Gaut1-1/gaut1-1* suspension cultured cells are smaller and have reduced HG:GalAT activity

The diminutive size of the stunted *gaut1-1/gaut1-1* seedlings was a major impediment in elucidating the effect(s) of loss of GAUT1 function on cell wall composition and architecture. We therefore used suspension cultured cells as an ample source of material for that purpose. Seeds from selfed *GAUT1/gaut1-1* were sown on callus-inducing medium, the genotype identities of the generated calli determined by DNA and transcript analyses, and each of the genotyped WT, *GAUT1/gaut1-1*, and *gaut1-1/gaut1-1* individual calli introduced into liquid medium to establish suspension cultures. *gaut1-1/gaut1-1* calli formed much more slowly (took several weeks longer to reach a sizable callus to inoculate liquid culture) than WT and *GAUT1/gaut1-1* calli, and were slightly darker in color as well as harder and less friable when manipulated (Figure S4A). This observation is in line with the absence of cell:cell adhesion defect in the *gaut1-1/gaut1-1* seedlings, and is opposite to the *qua1-1* mutant whose calli grew faster and were more friable than controls (Leboeuf et al., 2005). Observed under a scanning electron microscope, *gaut1-1/gaut1-1* calli were more extensively covered by fibrous extracellular material compared to WT calli (Figure S4B), suggesting that the *gaut1-1/gaut1-1* mutant calli secreted more extracellular material than WT cells and may explain the less friable characteristics of *gaut1-1/gaut1-1* calli.

The *gaut1-1/gaut1-1* suspension culture lines, once established, grew comparably to WT. The *gaut1-1/gaut1-1* suspension cultures had slightly lower dry weight of biomass accumulated over the first 10 days of culture but attained maximum biomass yield by day 12, suggesting a slower rate of cell division and/or cell expansion during the growth phase but production of slightly increased amounts of biomass compared to WT towards the end of the 14-day transfer cycle (Figure 5A). On a fresh weight basis, the *gaut1-1/gaut1-1* line had comparable biomass accumulation over the first 10 days of culture but significantly higher fresh weight biomass yield by day 12 (Figure S5). Consistent with the *gaut1-1/gaut1-1* mutation reduced cell expansion effect seen in seedlings, *gaut1-1/gaut1-1* suspension cultured cells were also significantly smaller in size compared to WT (Figure 5B-E). In line with the above observations on calli, the *gaut1-1/gaut1-1* suspension culture line secreted substantially larger amounts of extracellular material (ECM) into the culture medium, with 35.1 ± 3.7 (SD) mg ECM/100 mL culture for *gaut1-1/gaut1-1* compared to 14.6 ± 1.2 (SD) for WT (n = 3, *p* = 0.00598). These results are similar to the increased amount of extracellular material reported for *QUA1/GAUT8* mutant suspension culture (Leboeuf et al., 2005). Sugar composition analysis by gas chromatography-mass spectrometry (GC-MS) of trimethyl silyl glycosides of the ECM revealed relatively comparable amounts of most of the major (arabinose (Ara), xylose (Xyl), and glucose (Glc)) and minor (rhamnose (Rha), mannose (Man), fucose (Fuc), and glucuronic acid (GlcA)) sugar residues, but surprisingly showed significantly reduced galactose (Gal) content and more than three times higher GalA content in the *gaut1-1/gaut1-1* compared to WT samples (Figure S6). The latter suggests an increased secretion of HG into the medium of *gaut1-1/gaut1-1* suspension culture compared to WT. Since *gaut1-1/gaut1-1* was a knockout mutant line, we first confirmed the absence of GAUT1 at the protein level. Microsomal membrane preparations from WT, *GAUT1/gaut1-1*, and *gaut1-1/gaut1-1* suspension cell lines were separated by reducing and non-reducing gel electrophoresis, transferred to PVDF membranes, and probed with anti-GAUT1 polyclonal antibodies. The Western blot verified the absence of GAUT1 protein in the *gaut1-1/gaut1-1* line (Figure 5F). The microsomes were subsequently assayed for the incorporation of ^14^C-GalA from UDP-^14^C-GalA onto exogenous HG acceptor. Results of two independent assays of microsomes extracted from 7-day-old cells (harvested approximately 9 months and 11 months after the inception of the callus culture), showed a significant 22-27% reduction in HG:GalAT enzymatic activity in microsomes from *gaut1-1/gaut1-1* lines compared to WT (Figure 5G), consistent with GAUT1 biochemical function as an HG:GalAT. The remaining ∼75% of HG:GalAT activity in *gaut1-1/gaut1-1* suspension line microsomes was likely due to other HG:GalAT enzymes present in the microsomal membrane preparation. This is consistent with the relatively high transcript expression of other GAUTs (GAUTs 4, 8, 9, and 15) as revealed by quantitative PCR of the suspension culture cells (Figure S7). However, unlike in *gaut1-1/gaut1-1* seedlings, there seemed to be no transcript expression feedback response of other GAUTs in the *gaut1-1/gaut1-1* suspension culture line compared to WT.

**Figure 5.**
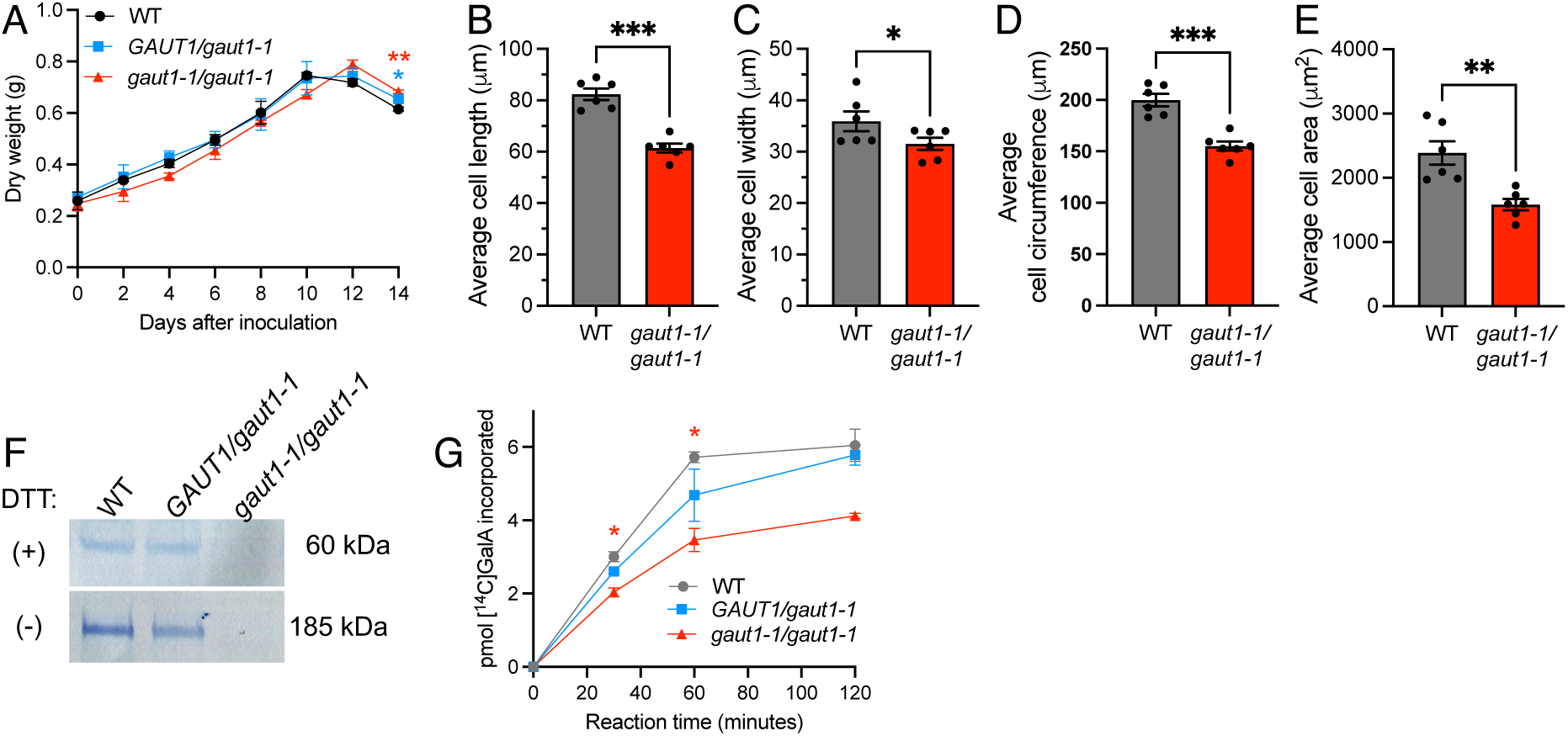
The *gaut1-1/gaut1-1* suspension cultured cell line had reduced growth, smaller cell size, and lower microsomal HG:GalAT activity compared to the WT line. (A) Growth of WT, *GAUT1/gaut1-1*, and *gaut1-1/gaut1-1* suspension cells over 14 days of culture, measured in dry weight. Data are means ± standard deviation from two independent experiments (n=2). Asterisks indicate significant difference to WT as analyzed by one-way Analysis of Variance (ANOVA) followed by Tukey multiple comparison test (* *P* < 0.05, ** *P* < 0.01). (B) Length, (C) width, (D) circumference, and (E) area of suspension cells of WT and *gaut1-1/gaut1-1* lines. Data are means ± standard error of duplicate measurements on three independent culture batches (n=6), with each data point represents the average of ≥ 100 cells. (F) Western blot of microsomes prepared from WT, *GAUT1/gaut1-1*, and *gaut1-1/gaut1-1* suspension cultured cells (harvested 9 to 11 months since inception of callus cultures) confirmed the absence of GAUT1 protein in the homozygous lines. Protein samples were separated by SDS-PAGE under reducing (+ DTT) or non-reducing (- DTT) condition and immunoblotted using an anti-GAUT1 antibody. (G) HG:GalAT activity in microsomes of 7-day-old WT, *GAUT1/gaut1-1* and *gaut1-1/gaut1-1* suspension cells (extracted from cells harvested 9 to 11 months since inception of callus cultures). Data are means ± standard deviation from a representative experiment performed in duplicates (n=2). Asterisks indicate significant difference compared to WT as determined by Student’s t-test (* *P* < 0.05, ** *P* < 0.01, *** *P* < 0.001).

During the course of this study we observed that the *gaut1-1/gaut1-1* suspension culture line appeared to adapt over time during successive subculturing by an increase in the overall total amount of HG:GalAT activity. We hypothesize that this is a type of compensation for the absence of GAUT1 that is associated with an increase in HG activity of one or more other GAUT HG:GalATs. Enzymatic assays of microsomes extracted from successively harvested cells over the 24-month period after the inception of the calli, revealed a progressive increase in total HG:GalAT activity in the *gaut1-1/gaut1-1* suspension culture lines which eventually surpassed the HG:GalAT activity of the WT control (Figure S13). The results highlight the importance of using calli or suspension culture cells early during the time period of culture initiation when generating material for cell wall analysis and mutant phenotype studies. The results also indicate the importance of monitoring biochemical characteristics of mutant cell cultures compared to WT controls to identify if biochemical properties change over time, potentially indicative of culture adaption.

### *Gaut1-1/gaut1-1* suspension cultured cells have reduced cell wall GalA, Man, cellulose, and calcium contents

To determine the effect of the GAUT1 mutation on cell wall composition and architecture, we isolated cell walls as Alcohol Insoluble Residues (AIRs) from WT, *GAUT1/gaut1-1* and *gaut1-1/gaut1-1* suspension cultured cell lines, removed starch by digestion with α-amylase, and performed sugar composition analysis (Figure 6A; Figure S8). We found that the total wall GalA content in *gaut1-1/gaut1-1* cells was significantly 30% reduced compared to that of heterozygous and WT cells (Figure 6A), indicating reduced HG content in the absence of GAUT1. The total wall Man content was significantly lower in *gaut1-1/gaut1-1* compared to WT, suggesting that the production of mannans may also be affected by the *gaut1* mutation. On the other hand, Ara content was significantly increased, likely as a compensatory response to GAUT1’s absence. Furthermore, the cellulose content of the AIR samples was markedly lower in the homozygous *gaut1-1/gaut1-1* mutant compared to the heterozygous and WT lines (Figure 6B).

**Figure 6.**
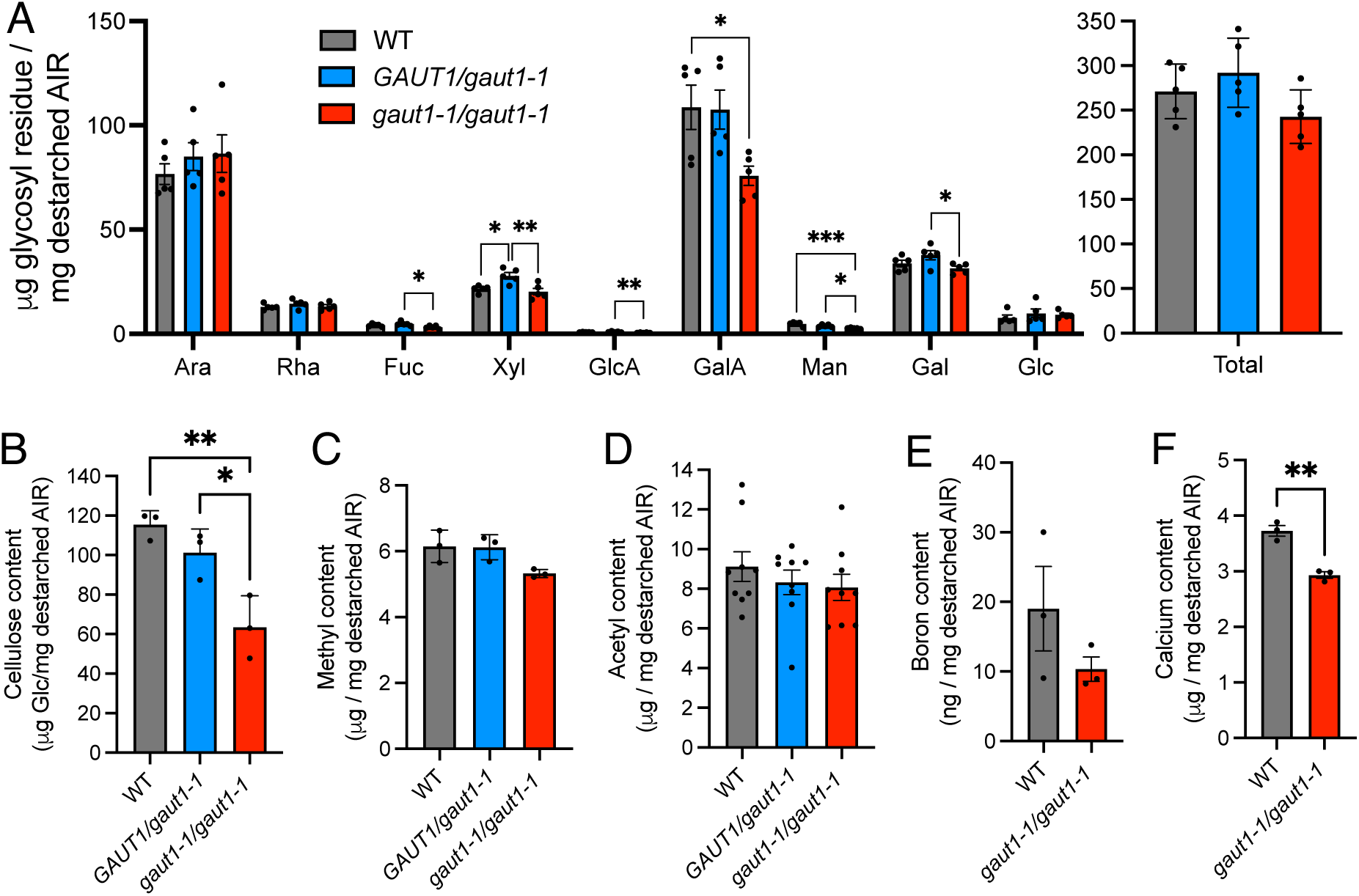
Analyses of total cell walls revealed lower GalA content in the *gaut1-1/gaut1-1* compared to WT and *GAUT1/gaut1-1* suspension culture lines. (A) Individual and total non-cellulosic glycosyl residue content of destarched total cell walls (alcohol insoluble residue (AIR)), presented in mass (μg/mg destarched AIR), from WT, *GAUT1/gaut1-1* and *gaut1-1/gaut1-1* suspension cultures (log phase growth), as measured by GC-MS of trimethylsilyl (TMS) derivatives. Data are means ± standard error of two technical replicates of destarched AIR extracted from five independent culture batches (harvested ∼1 to 2.5 years after inception of callus cultures). (B) Cellulose, (C) methyl, (D) acetyl, (E) boron, and (F) calcium content of the destarched total cell wall from WT, *GAUT1/gaut1-1*, and *gaut1-1/gaut1-1* suspension culture lines. Data are means ± standard error of three independent culture batches. Asterisks indicate significant difference as determined by one-way ANOVA followed by Tukey’s multiple comparison test for panels (A) and (B), and by Student’s t-test for panel (F) (* *P* < 0.05, ** *P* < 0.01, *** *P* < 0.001).

The GalA residues in pectin may be methylesterified and/or acetylated, and pectic polymers may be crosslinked with each other via HG-calcium crosslinking and/or borate-diester dimerization of RG-II. Therefore, the level of these modifications and the elemental content of total cell walls were measured. There was a trend toward lower methyl, acetyl, and boron content in the total walls from *gaut1-1/gaut1-1* compared to heterozygous and WT lines; however, these trends were not statistically significant (Figure 6C-E). In line with the reduced amount of GalA/HG content, the calcium content of the total walls was significantly lower in the *gaut1-1/gaut1-1* cell line compared to WT (Figure 6F). Taken together, the data support the conclusion that HG-calcium crosslinking is reduced in *gaut1-1/gaut1-1* walls.

### Glycosyl residue and linkage compositions of cell wall fractions from suspension culture cells reveal significantly reduced GalA in the most tightly bound fractions of *gaut1-1/gaut1-1* walls

GAUT1 belongs to the *GAUT* gene family with 15 members in Arabidopsis. Several of these members, including GAUT1, have been biochemically demonstrated to possess HG:GalAT activity (Sterling et al., 2006; Biswal et al., 2018b; Voiniciuc et al., 2018; Engle et al., 2022), leading us to hypothesize that the different GAUTs synthesize HG glycans that may stand alone or be part of more complex wall polymers (Atmodjo et al., 2013). To test this hypothesis for GAUT1, we performed six sequential extractions of cell walls from WT, *GAUT1/gaut1-1*, and *gaut1-1/gaut1-1* suspension culture lines using increasingly harsh solvents (see below and Materials and Methods) and analyzed the wall extracts obtained. Significantly more material was extracted from *gaut1-1/gaut1-1* cell walls in the last/harshest 4M KOH post-chlorite solvent extraction step (82% more) compared to the same extract sample from WT cell walls (Figure S9A), suggesting increased extractability of tightly-bound wall polymers when GAUT1-synthesized HG is not produced.

The wall extracts and insoluble residue from *gaut1-1/gaut1-1* and WT were then analyzed by glycosyl residue composition to determine if wall sugar composition was altered due to the GAUT1 mutation, and if so, whether specific wall fractions were affected. Table 2 shows glycosyl residue composition as mass yield recovered in each wall fraction (µg sugar/mg de-starched AIR). The first two wall fractions, extracted sequentially by ammonium oxalate and sodium carbonate to remove polymers held in the wall respectively by electrostatic interactions and ester linkages, are generally enriched in pectic polysaccharides. The next two fractions, 1M KOH and 4M KOH extracts, are expected to be enriched in hemicelluloses. The 1M KOH extracts glycans held in the wall by hydrogen bonds and/or by hydrolyzable ester linkages between glycan and hydroxycinnamic acid. The 4M KOH extracts glycans held more tightly in the wall by hydrogen bonds. Subsequent treatment with sodium chlorite/acetic acid is typically used to delignify tissues with secondary cell walls such as lignocellulosic biomass. For non-lignified tissues such treatment can be expected to break phenolic and phenolic-ester cross-linkages to polysaccharides (Selvendran, 1985). Finally, a second 4M KOH extraction step after the sodium chlorite/acetic acid treatment extracts polymers whose connection to the wall matrix was cleaved by the sodium chlorite/acetic acid but then become releasable by base extraction, e.g., polymers held in the wall by hydrogen bonding.

**Table 2.**
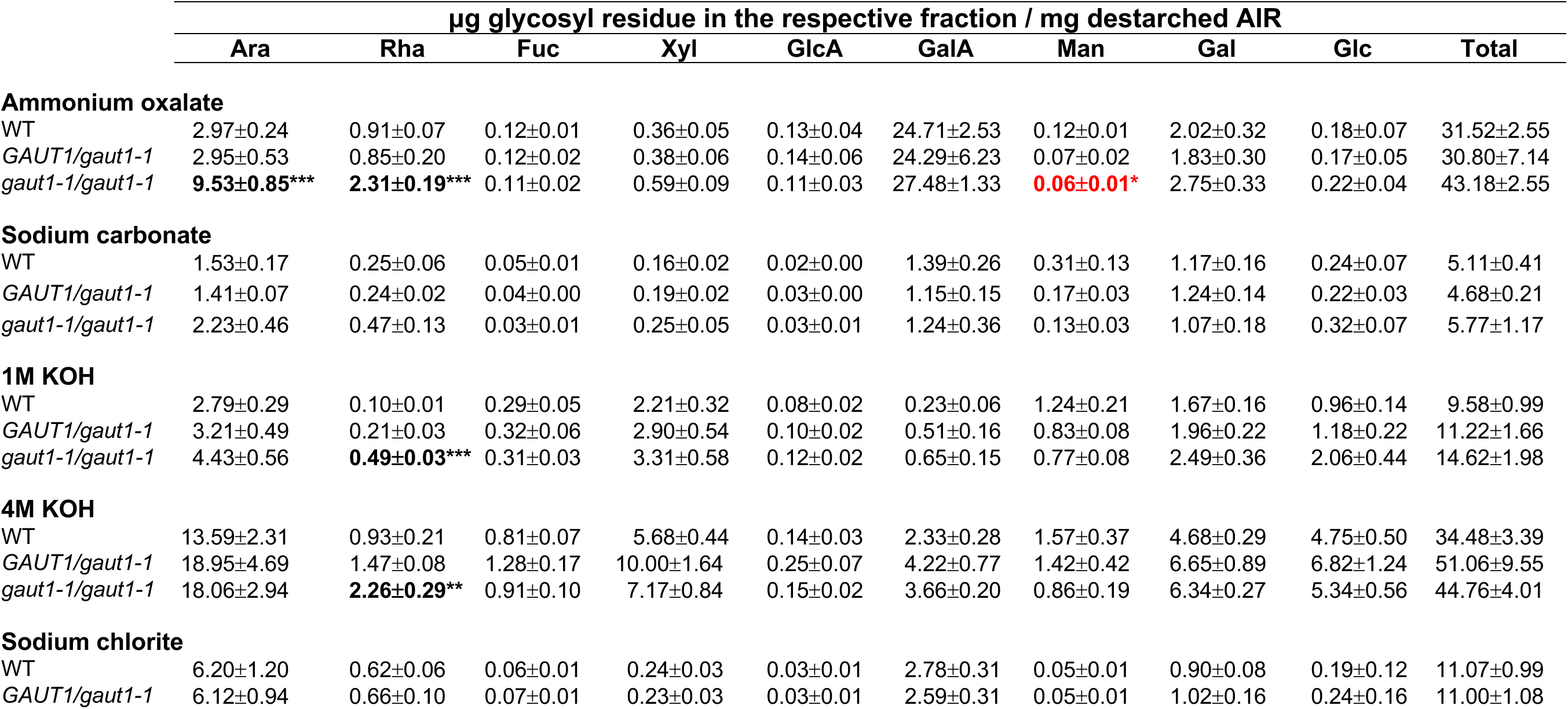
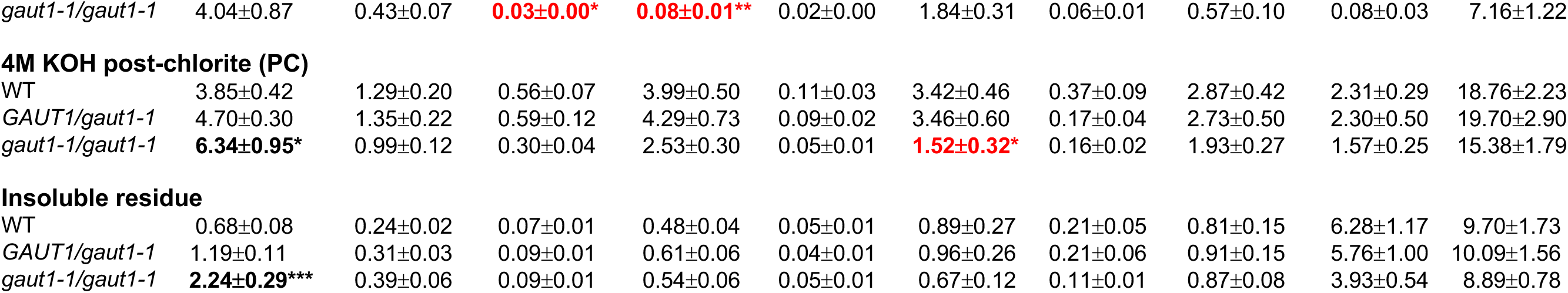
Glycosyl residue composition of cell wall fractions of WT, *GAUT1/gaut1-1*, and *gaut1-1/gaut1-1* suspension culture cell lines. Destarched AIR was extracted sequentially using 50 mM ammonium oxalate, 50 mM sodium carbonate, 1M KOH, 4M KOH, 100 mM sodium chlorite, and 4M KOH post-chlorite (PC), and the resulting fractions as well as the insoluble residue were analyzed by GC-MS of trimethylsilyl (TMS) derivatives. Data are means ± standard error of two technical replicates of wall fractions from five independent culture batches (n = 5). Asterisks represent statistically significant difference in comparison to WT at \**P* < 0.05, \*\**P* < 0.01, and \*\*\**P* < 0.001 (one-way ANOVA followed by Tukey’s multiple comparison test). Red denotes statistically significant reduction compared to WT.

|  | **$\mu\text{g}$ glycosyl residue in the respective fraction / mg destarched AIR** |  |  |  |  |  |  |  |  |  |
| --- | --- | --- | --- | --- | --- | --- | --- | --- | --- | --- |
|  | **Ara** | **Rha** | **Fuc** | **Xyl** | **GlcA** | **GalA** | **Man** | **Gal** | **Glc** | **Total** |
| **Ammonium oxalate** |  |  |  |  |  |  |  |  |  |  |
| WT | 2.97±0.24 | 0.91±0.07 | 0.12±0.01 | 0.36±0.05 | 0.13±0.04 | 24.71±2.53 | 0.12±0.01 | 2.02±0.32 | 0.18±0.07 | 31.52±2.55 |
| *GAUT1/gaut1-1* | 2.95±0.53 | 0.85±0.20 | 0.12±0.02 | 0.38±0.06 | 0.14±0.06 | 24.29±6.23 | 0.07±0.02 | 1.83±0.30 | 0.17±0.05 | 30.80±7.14 |
| *gaut1-1/gaut1-1* | **9.53±0.85\*\*\*** | **2.31±0.19\*\*\*** | 0.11±0.02 | 0.59±0.09 | 0.11±0.03 | 27.48±1.33 | **0.06±0.01\*** | 2.75±0.33 | 0.22±0.04 | 43.18±2.55 |
| **Sodium carbonate** |  |  |  |  |  |  |  |  |  |  |
| WT | 1.53±0.17 | 0.25±0.06 | 0.05±0.01 | 0.16±0.02 | 0.02±0.00 | 1.39±0.26 | 0.31±0.13 | 1.17±0.16 | 0.24±0.07 | 5.11±0.41 |
| *GAUT1/gaut1-1* | 1.41±0.07 | 0.24±0.02 | 0.04±0.00 | 0.19±0.02 | 0.03±0.00 | 1.15±0.15 | 0.17±0.03 | 1.24±0.14 | 0.22±0.03 | 4.68±0.21 |
| *gaut1-1/gaut1-1* | 2.23±0.46 | 0.47±0.13 | 0.03±0.01 | 0.25±0.05 | 0.03±0.01 | 1.24±0.36 | 0.13±0.03 | 1.07±0.18 | 0.32±0.07 | 5.77±1.17 |
| **1M KOH** |  |  |  |  |  |  |  |  |  |  |
| WT | 2.79±0.29 | 0.10±0.01 | 0.29±0.05 | 2.21±0.32 | 0.08±0.02 | 0.23±0.06 | 1.24±0.21 | 1.67±0.16 | 0.96±0.14 | 9.58±0.99 |
| *GAUT1/gaut1-1* | 3.21±0.49 | 0.21±0.03 | 0.32±0.06 | 2.90±0.54 | 0.10±0.02 | 0.51±0.16 | 0.83±0.08 | 1.96±0.22 | 1.18±0.22 | 11.22±1.66 |
| *gaut1-1/gaut1-1* | 4.43±0.56 | **0.49±0.03\*\*\*** | 0.31±0.03 | 3.31±0.58 | 0.12±0.02 | 0.65±0.15 | 0.77±0.08 | 2.49±0.36 | 2.06±0.44 | 14.62±1.98 |
| **4M KOH** |  |  |  |  |  |  |  |  |  |  |
| WT | 13.59±2.31 | 0.93±0.21 | 0.81±0.07 | 5.68±0.44 | 0.14±0.03 | 2.33±0.28 | 1.57±0.37 | 4.68±0.29 | 4.75±0.50 | 34.48±3.39 |
| *GAUT1/gaut1-1* | 18.95±4.69 | 1.47±0.08 | 1.28±0.17 | 10.00±1.64 | 0.25±0.07 | 4.22±0.77 | 1.42±0.42 | 6.65±0.89 | 6.82±1.24 | 51.06±9.55 |
| *gaut1-1/gaut1-1* | 18.06±2.94 | **2.26±0.29\*\*** | 0.91±0.10 | 7.17±0.84 | 0.15±0.02 | 3.66±0.20 | 0.86±0.19 | 6.34±0.27 | 5.34±0.56 | 44.76±4.01 |
| **Sodium chlorite** |  |  |  |  |  |  |  |  |  |  |
| WT | 6.20±1.20 | 0.62±0.06 | 0.06±0.01 | 0.24±0.03 | 0.03±0.01 | 2.78±0.31 | 0.05±0.01 | 0.90±0.08 | 0.19±0.12 | 11.07±0.99 |
| *GAUT1/gaut1-1* | 6.12±0.94 | 0.66±0.10 | 0.07±0.01 | 0.23±0.03 | 0.03±0.01 | 2.59±0.31 | 0.05±0.01 | 1.02±0.16 | 0.24±0.16 | 11.00±1.08 |
| *gaut1-1/gaut1-1* | 4.04±0.87 | 0.43±0.07 | 0.03±0.00* | 0.08±0.01** | 0.02±0.00 | 1.84±0.31 | 0.06±0.01 | 0.57±0.10 | 0.08±0.03 | 7.16±1.22 |
| **4M KOH post-chlorite (PC)** |  |  |  |  |  |  |  |  |  |  |
| WT | 3.85±0.42 | 1.29±0.20 | 0.56±0.07 | 3.99±0.50 | 0.11±0.03 | 3.42±0.46 | 0.37±0.09 | 2.87±0.42 | 2.31±0.29 | 18.76±2.23 |
| *GAUT1/gaut1-1* | 4.70±0.30 | 1.35±0.22 | 0.59±0.12 | 4.29±0.73 | 0.09±0.02 | 3.46±0.60 | 0.17±0.04 | 2.73±0.50 | 2.30±0.50 | 19.70±2.90 |
| *gaut1-1/gaut1-1* | 6.34±0.95* | 0.99±0.12 | 0.30±0.04 | 2.53±0.30 | 0.05±0.01 | 1.52±0.32* | 0.16±0.02 | 1.93±0.27 | 1.57±0.25 | 15.38±1.79 |
| **Insoluble residue** |  |  |  |  |  |  |  |  |  |  |
| WT | 0.68±0.08 | 0.24±0.02 | 0.07±0.01 | 0.48±0.04 | 0.05±0.01 | 0.89±0.27 | 0.21±0.05 | 0.81±0.15 | 6.28±1.17 | 9.70±1.73 |
| *GAUT1/gaut1-1* | 1.19±0.11 | 0.31±0.03 | 0.09±0.01 | 0.61±0.06 | 0.04±0.01 | 0.96±0.26 | 0.21±0.06 | 0.91±0.15 | 5.76±1.00 | 10.09±1.56 |
| *gaut1-1/gaut1-1* | 2.24±0.29*** | 0.39±0.06 | 0.09±0.01 | 0.54±0.06 | 0.05±0.01 | 0.67±0.12 | 0.11±0.01 | 0.87±0.08 | 3.93±0.54 | 8.89±0.78 |

Since GAUT1 is an HG:GalAT, we hypothesized that the absence of GAUT1 would cause a reduction in GalA content in one or more wall fractions, and that those comparable fractions in WT should be sources of GAUT1-synthesized HG. Sugar composition analysis (µg sugar/mg destarched AIR) of different wall fractions of *gaut1-1/gaut1-1* versus WT (Table 2), revealed a significant reduction of GalA compared to WT only in the 4M KOH post-chlorite fraction (56%). Significant reductions were also observed for Man in the ammonium oxalate fraction and for Fuc and Xyl in the sodium chlorite fractions of *gaut1-1/gaut1-1* samples compared to WT, while Ara and Rha contents were increased significantly in several wall extracts. Notably, the substantial reduction in GalA content in only the 4M KOH post-chlorite fraction in the *gaut1-1/gaut1-1* mutant was different than the location of reduced GalA content previously reported for transgenic plants down-regulated in GAUT4 (Biswal et al., 2018b) and GAUT12 (Biswal et al., 2015), where the GalA content was found consistently to be significantly reduced in all six wall extracts and in the remaining insoluble residue. Thus, the results support the hypothesis that GAUT1 synthesizes a specific type of HG that in WT is largely isolated in the 4M KOH post-chlorite fraction from WT walls.

We further determined the glycosyl linkage composition of four wall fractions that showed reduced or comparable GalA content in the *gaut1-1/gaut1-1* line compared to WT, i.e. ammonium oxalate, sodium carbonate, sodium chlorite, and 4M KOH post-chlorite fractions (Table 3). The ammonium oxalate fractions showed a >40% reduction in 3,4-linked GalA and terminal-Xyl, glycosyl linkages typically present in xylogalacturonan, in *gaut1-1/gaut1-1* compared to WT samples. In line with the increased Ara and Rha content in the ammonium oxalate fractions, linkage analysis showed a marked increase in multiple Ara and Rha linkage types including t-Ara*f*, 4- or 5-Ara*p*, 2,4-Ara*p*, 2,3,4-Ara*p*, 2-Rha and 2,4-Rha in *gaut1-1/gaut1-1* compared to WT, suggesting a compensatory increase of arabinan and RG-I or other Rha-containing polysaccharides in this fraction in the absence of GAUT1-synthesized HG. Consistent with these trends, boron, calcium, and methyl content of the ammonium oxalate fractions were comparable while acetyl content, presumably associated with RG-I backbone, was increased in the *gaut1-1/gaut1-1* ammonium oxalate fraction compared to WT (Figure S9 B-E). The sodium carbonate fractions had reduced amounts of terminal-linked, 4-linked, and 3,4-linked GalAs in *gaut1-1/gaut1-1* compared to WT, consistent with reduction in HG and possibly xylogalacturonan (Table 3). Since the methyl content was significantly reduced while calcium content was increased (Figure S9 C-D), it is plausible that the HG remaining in the sodium carbonate fraction of *gaut1-1/gaut1-1* is less methylesterified and thus engaged more in calcium crosslinking than in the WT samples. Interestingly, in the sodium chlorite and 4M KOH post-chlorite wall fractions only 4-GalA was reduced in *gaut1-1/gaut1-1* in comparison to WT (Table 3), in support of HG being reduced in these two final wall fractions especially in the 4M KOH post-chlorite. Furthermore, multiple Rha-linked species and several Gal-linked species were also lower in *gaut1-1/gaut1-1* than WT samples for these fractions, which are possibly attributed to RG-I backbone and galactan side chains, while many Ara-linked species were upregulated suggesting a compensatory increase in arabinan due to GAUT1 mutation. In keeping with the significantly reduced amount of Man in the ammonium oxalate fraction of *gaut1-1/gaut1-1* in comparison to WT (Tables 2), the glycosyl linkage analysis revealed corresponding decreases in t-Man and 4-Man (Table 3), suggesting a possible type of association of mannan to the GAUT1-synthesized HG.

**Table 3.** Glycosyl linkage composition of ammonium oxalate, sodium carbonate, sodium chlorite, and 4M KOH post-chlorite wall fractions extracted from WT and *gaut1-1/gaut1-1* suspension cultured cells. Data are in mol percentages.

|  | Ammonium oxalate |  | Sodium carbonate |  | Sodium chlorite |  | 4M KOH-PC |  |
| --- | --- | --- | --- | --- | --- | --- | --- | --- |
|  | WT | *gaut1-1/gaut1-1* | WT | *gaut1-1/gaut1-1* | WT | *gaut1-1/gaut1-1* | WT | *gaut1-1/gaut1-1* |
| t-Araf | 11.8 | 22.1 | 13.75 | 16.0 | 14.2 | 15.4 | 6.0 | 9.7 |
| t-Arap | - | 0.3 | 0.35 | 0.5 | 0.7 | 1.0 | 0.3 | 0.5 |
| 2-Araf | 1.2 | 0.8 | 1.65 | 1.0 | 4.6 | 3.7 | 2.0 | 1.4 |
| 3-Araf | 2.2 | 1.9 | 2.3 | 1.5 | 3.5 | 4.7 | 1.3 | 1.8 |
| 4-Arap or 5-Araf | 7.1 | 12.2 | 5.6 | 7.8 | 6.7 | 8.1 | 3.6 | 4.9 |
| 2,4-Arap or 2,5-Araf | 2.5 | 4.6 | 1.95 | 3.2 | 1.9 | 3.0 | 1.0 | 2.1 |
| 2,3,4-Arap or 2,3,5-Araf | 2.8 | 6.2 | 2.3 | 4.4 | 2.8 | 4.0 | 1.5 | 3.6 |
| 2-Rhap | 1.3 | 2.7 | 1.55 | 1.8 | 2.4 | 2.2 | 1.9 | 1.6 |
| 4-Rhap | 0.6 | 0.4 | 0.2 | 0.3 | 0.6 | 0.5 | 0.3 | 0.3 |
| 2,4-Rhap | 1.7 | 3.4 | 2.6 | 2.6 | 4.2 | 3.7 | 4.0 | 3.3 |
| t-Fuc | 1.2 | 0.4 | 0.65 | 0.7 | 1.4 | 0.8 | 3.1 | 1.9 |
| t-Xylp | 2.9 | 1.4 | 2.65 | 2.4 | 2.5 | 1.9 | 7.4 | 6.6 |
| 2-Xylp | 0.6 | 0.3 | 0.7 | 0.7 | 0.3 | 0.5 | 2.1 | 1.7 |
| 4-Xylp | 3.0 | 1.1 | 3.9 | 1.8 | 2.6 | 3.0 | 3.5 | 2.2 |
| 2,4-Xylp | 0.9 | - | 0.35 | 0.6 | 0.3 | 0.2 | 1.4 | 0.4 |
| t-GlcAp | 1.5 | 0.7 | 1.05 | 0.7 | 0.7 | 0.7 | 0.9 | 0.5 |
| 2-GlcAp | 0.8 | 0.6 | 0.5 | 0.5 | 1.3 | 0.8 | 1.8 | 0.8 |
| t-GalAp | 1.0 | 1.1 | 0.7 | 0.4 | 0.7 | 0.6 | 0.3 | 0.4 |
| 2-GalAp | 0.3 | 0.2 | 0.1 | 0.3 | 0.1 | 0.2 | - | - |
| 4-GalAp | 7.0 | 10.4 | 5.95 | 4.3 | 10.7 | 7.8 | 6.4 | 5.7 |
| 3,4-GalAp | 2.8 | 1.6 | 2.25 | 1.5 | 2.2 | 2.8 | 0.9 | 1.0 |
| t-Manp | 0.5 | 0.2 | 0.8 | 0.5 | 0.2 | 0.5 | 0.2 | 0.2 |
| 4-Manp | 3.9 | 1.6 | 5.1 | 2.8 | 2.9 | 3.4 | 3.8 | 2.5 |
| 4,6-Manp | - | 3.4 | 1.7 | 1.6 | 0.3 | - | 2.4 | 1.1 |
| t-Galp | 3.0 | 2.8 | 4.05 | 4.0 | 3.4 | 3.2 | 4.3 | 4.5 |
| 2-Galp | 0.7 | 0.4 | 0.7 | 0.5 | 0.2 | 0.2 | 3.4 | 2.5 |
| 3-Galp | 2.6 | 1.5 | 2.05 | 2.1 | 2.0 | 1.6 | 1.8 | 2.3 |
| 4-Galp | 1.0 | 4.1 | 2.45 | 2.7 | 3.5 | 3.7 | 3.8 | 3.4 |
| 6-Galp | 2.9 | 1.2 | 2.8 | 2.0 | 1.8 | 1.6 | 1.1 | 1.2 |
| 3,4-Galp | 2.1 | 0.6 | 1.6 | 1.3 | 1.4 | 1.8 | 0.6 | 0.8 |
| 3,6-Galp | 9.2 | 3.1 | 8.75 | 6.0 | 2.9 | 2.8 | 1.4 | 1.9 |
| 4,6-Galp | - | 0.1 | 0.2 | 0.3 | 0.1 | 0.1 | 0.1 | 0.2 |
| t-Glcp | 1.9 | 0.5 | 0.75 | 1.2 | 0.7 | 1.0 | 0.5 | 0.9 |
| 3-Glcp | 1.4 | 0.5 | 6.85 | 10.8 | 0.4 | 0.7 | 1.1 | 1.6 |
| 2-Glcp | 0.3 | 0.2 | 0.1 | 0.3 | 0.2 | 0.1 | 0.1 | 0.2 |
| 4-Glcp | 15.7 | 7.2 | 8.9 | 9.4 | 13.7 | 12.2 | 12.0 | 13.1 |
| 4,6-Glcp | 3.6 | 2.8 | 2.4 | 3.5 | 2.4 | 2.5 | 13.7 | 14.7 |

### Glycome profiling analysis shows altered extraction patterns of xyloglucan, xylan, HG, and galactan epitopes in *gaut1-1/gaut1-1* suspension cultured cells

We also analyzed the sequentially extracted wall fractions by glycome profiling, an ELISA-based assay using a series of antibodies directed against various plant cell wall glycans (Pattathil et al., 2010) to detect different cell wall glycan epitopes released upon fractionation of the wall with different solvents (Pattathil et al., 2012). We used the same sequence of reagents to prepare the cell wall extracts as mentioned above to prepare the sequential wall extracts. The ELISA signals for 77 monoclonal antibodies against the six wall extracts of three independently grown suspension cultures of WT and *gaut1-1/gaut1-1* lines were measured (Figure 7).

**Figure 7.**
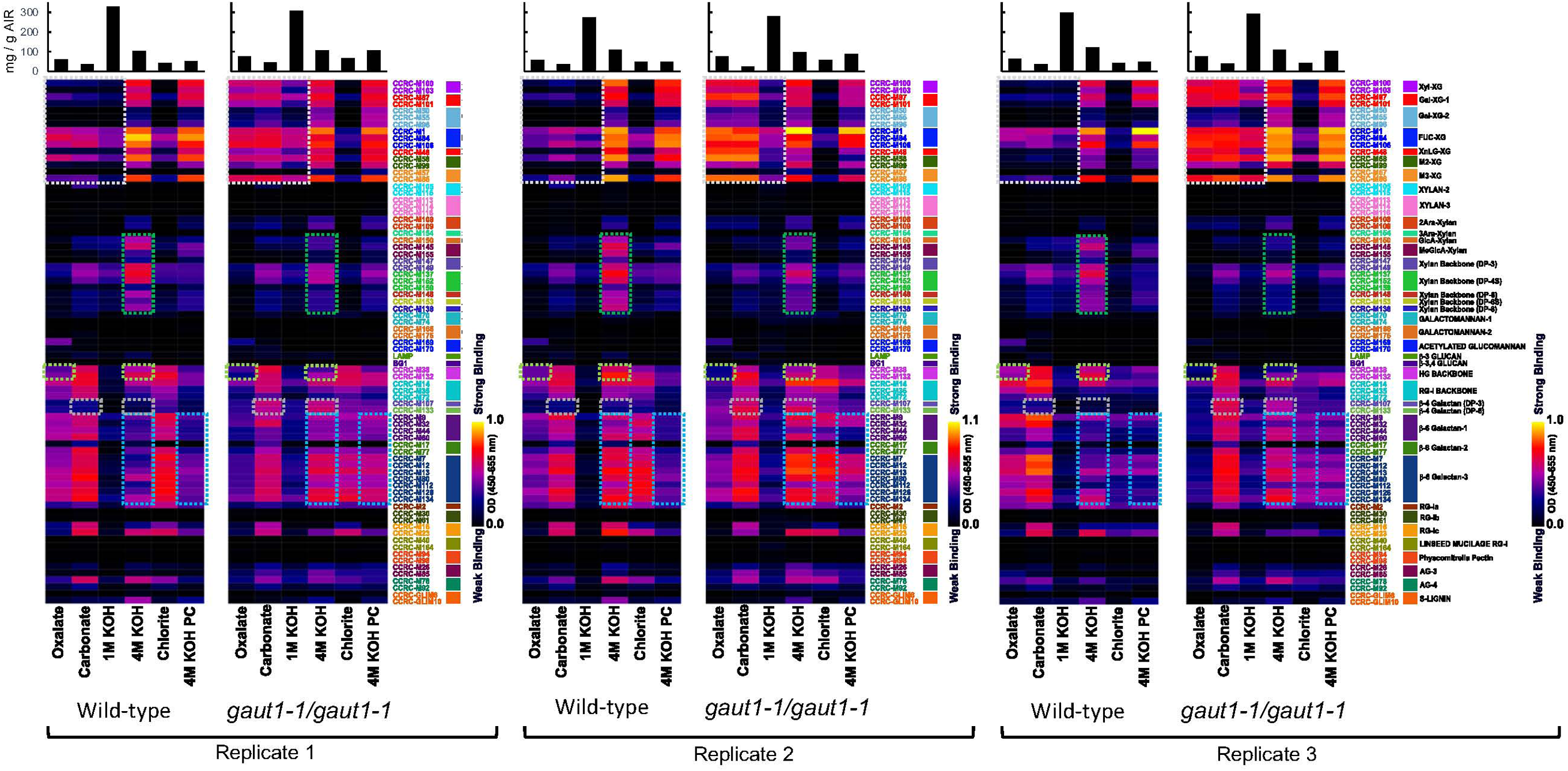
Glycome profiling of WT and *gaut1-1/gaut1-1* suspension cultured cells. Six wall fractions were sequentially extracted from respective cell wall (AIR) preparations from three independently grown cell cultures (harvested ∼2 to 2.5 years after inception of callus cultures), using increasingly harsh solvents as follows: 50 mM ammonium oxalate (Oxalate), 50 mM sodium carbonate (Carbonate), 1M KOH, 4M KOH, acidified sodium chlorite (Chlorite), and 4M KOH post-chlorite (4M KOH PC). The resulting wall fractions were screened for glycan epitope content via ELISA using a suite of 77 monoclonal antibodies (mAbs) directed against plant cell wall glycan epitopes. The antibody binding strength is represented as heatmaps using a white, red, and dark blue scale (depicting strong, medium, and no binding, respectively). Bar graphs above the heatmaps show the actual amounts of cell wall material extracted in each extraction step. Color-coded panels to the right of heatmaps show the antibodies used, which are grouped based on the main cell wall glycans they recognize. Observed major changes (increase or decrease) in the antibody binding response to mutant versus WT wall fractions were outlined in dotted boxes (white – xyloglucan epitopes, dark green – xylan epitopes, light green – HG backbone epitopes, grey – β-1,4-galactan epitopes, and light blue – β-1,6-galactan epitopes).

Several consistent changes in the pattern of release of specific types of cell wall polymers from the wall were observed across the three replicates of the glycome profiles of the *gaut1-1/gaut1-1* mutant compared to WT. First and foremost, recognition signals of xyloglucan epitopes by multiple anti-xyloglucan antibodies increased markedly in the early released wall fractions of walls isolated from the *gaut1-1/gaut1-1-* mutant compared to WT, namely in the Oxalate and Carbonate fractions, and to a lesser extent in the 1M KOH fractions (Figure 7, white boxes). This is a significant change considering that xyloglucans are not typically released in these early wall fractions but instead are generally extracted from the wall predominantly by strong base treatment (4M KOH or NaOH) (Park and Cosgrove, 2015). Such early release of xyloglucans from the *gaut1-1/gaut1-1* mutant walls indicates that the loss of GAUT1-synthesized HG causes substantial wall architectural change that results in xyloglucans not being held in the wall as tightly as they normally are in WT walls. Secondly, for the xylan-reacting antibodies there was a reduction in the signal strength for recognition of both 4-O-methylglucuronoxylan and xylan backbone epitopes, particularly in the 4M KOH fractions of *gaut1-1/gaut1-1* mutant compared to WT (Figure 7, dark green boxes), suggesting that loss of GAUT1-synthesized HG causes a reduction in the amounts of xylans in these fractions of the mutant walls.

Thirdly, there was markedly reduced binding of anti-HG backbone antibodies CCRC-M38 and CCRC-M132 in the Oxalate and 4M KOH fractions of the *gaut1-1/gaut1-1* mutant compared to WT (Figure 7, light green boxes). This result indicates that at least some GAUT1-synthesized HG is held in the walls by ionic interactions (releasable by ammonium oxalate) while other GAUT1-synthesized HG is tightly held into the wall (releasable by the harsher solvent 4M KOH). In contrast to HG backbone, there were no reproducible pattern shifts in immunoreactivity against RG-I backbone epitopes, despite a slight tendency of reduced anti-RG-I backbone antibody binding in different cell wall fractions of *gaut1-1/gaut1-1* mutant compared to WT. Finally, there was increased binding signal strength for anti-β-1,4-galactan antibodies CCRC-M107 and CCRC-M133 in the Carbonate and 4M KOH fractions (Figure 7, grey boxes), and for multiple anti-β-1,6-galactan antibodies in the 4M KOH and 4M KOH PC fractions (Figure 7, light blue boxes) of *gaut1-1/gaut1-1* cell walls compared to WT. Taken together, the glycome profiling results indicate that cell wall structure and architecture is loosened in Arabidopsis GAUT1 mutant suspension culture cells, leading to more easily extracted xyloglucans and galactans. Reduced HG in the ammonium oxalate and 4 M KOH fractions was associated with loss of GAUT1-synthesized HG. The lack of effect on RG-I backbone signal but increased signal for β-1,4- and β-1,6-galactans suggest that while overall RG-I synthesis did not appear upregulated in the mutant, the amount of side-branching was increased.

### GAUT1-synthesized HG in WT is connected to an RG-II glycan band that is downregulated in *gaut1-1/gaut1-1* suspension culture cell walls

Differential extraction of cell walls with solvents of increasing ionic strength and chemical reactivity leads to separation of wall polysaccharides and proteoglycans held in the wall by different types of ionic and covalent bonds. To determine if there were differences in the size and/or distribution of negatively charged wall polymers, likely pectic polymers, in the different wall fractions of WT and *gaut1-1/gaut1-1* suspension cells, aliquots of each wall fraction were analyzed by high percentage polyacrylamide gel electrophoresis (HP-PAGE) followed by staining with alcian blue and silver nitrate (Amos et al., 2018). This gel method detects negatively charged PAGE-separated glycans by staining with the positively charged stain alcian blue followed by silver stain to fix the signal in the gels. Direct loading of equal amounts of each wall fraction onto the gel followed by electrophoresis resulted in a smear and some banding patterns as shown in Figure 8A. The strongest staining, taken to represent an abundance of negatively charged pectins, was observed in the ammonium oxalate and sodium carbonate fractions that are known to be enriched in pectins. These two fractions also showed the presence of ladder-like bands of oligogalacturonides (HG oligomers) in the bottom half of the lanes. Relatively strong staining was also observed in the smears in the upper part of the sodium chlorite and 4M KOH post-chlorite fractions. The 1M KOH and 4M KOH fractions had the lightest staining, consistent with these fractions being hemicellulose-enriched. Differences in the smear/band patterns were observed between the WT and *gaut1-1/gaut1-1* samples. These included (1) different band patterns in the upper part of the ammonium oxalate fraction; (2) stronger staining in the WT compared to *gaut1-1/gaut1-1* in the upper part of the 1M and 4M KOH fractions; and (3) the presence of a faint but distinct band in the middle of the lane for WT and a much darker smear in the top half of the lane for *gaut1-1/gaut1-1* of the 4M KOH post-chlorite fraction.

**Figure 8.**
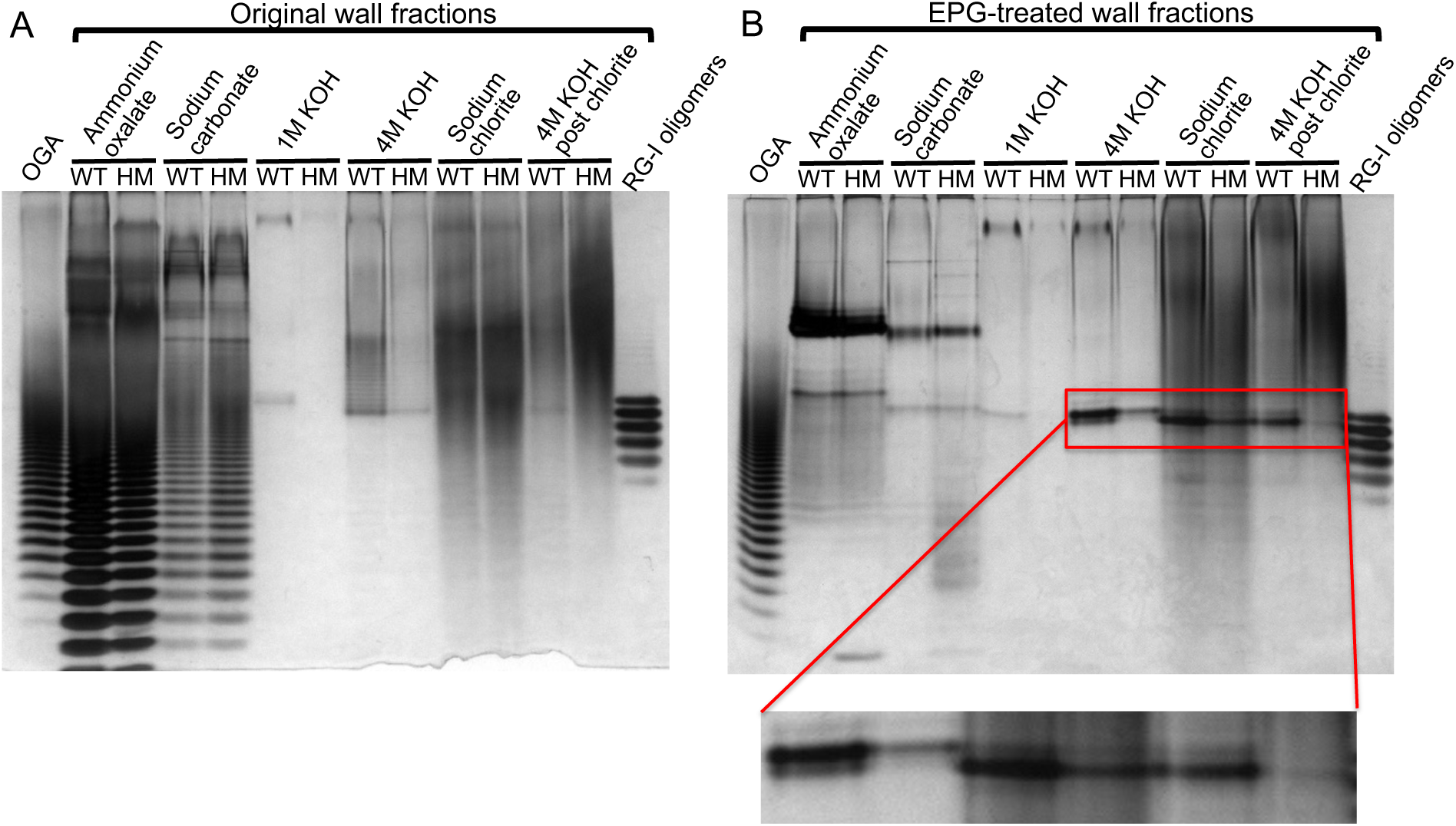
High percentage polyacrylamide gel electrophoresis (HP-PAGE) of wall fractions of WT and *gaut1-1/gaut1-1* suspension culture lines, without (A) and with (B) pre-treatment with endopolygalacturonase (EPG). Data are representative results from at least 3 independent culture batches. HM – homozygous *gaut1-1/gaut1-1* mutant. Inset: enlarged picture of the distinct doublet polymer bands in 4M KOH, sodium chlorite, and 4M KOH post-chlorite fractions, that were downregulated in *gaut1-1/gaut1-1* samples compared to WT.

To identify the polymer that was HG, the wall fractions were treated with endopolygalacturonase (EPG), which cleaves HG, and the hydrolysates were then analyzed by gel electrophoresis (Figure 8B). As expected, the EPG treatment abolished the oligogalacturonide ladder-like bands and most of the upper smears in the ammonium oxalate and sodium carbonate fractions, confirming that those materials were HG. We also observed, in the EPG-treated ammonium oxalate and sodium carbonate fractions, two major bands in the upper half of the lanes at the approximate position where RG-II dimer and monomer, respectively, electrophorese during HP-PAGE (Figures 8B and S10). The intensity of the bands was roughly comparable between the WT and the *gaut1-1/gaut1-1* wall extracts, suggesting that the bulk of RG-II extracted by ammonium oxalate and sodium carbonate is not substantially affected by the *gaut1-1/gaut1-1* mutation. We noted that the two major bands in the sodium carbonate fractions migrated faster during HP-PAGE compared to those in the ammonium oxalate fractions. This is likely due to removal of ester bonds from the RG-II backbone and/or side chains upon sodium carbonate extraction, resulting in more anionic and smaller RG-II monomer and dimer that moved faster towards the positive electrode during electrophoresis. No change in the band pattern or intensity was observed in WT and *gaut1-1/gaut1-1* EPG-pretreated 1M KOH fractions analyzed by HP-PAGE. However, the EPG pre-treatment of the 4M KOH, sodium chlorite, and 4M KOH post-chlorite fractions revealed the appearance and/or darkening of distinct doublet of polymer bands (see inset in Figure 8B) at approximately the midpoint of the lanes in all three wall extracts, with a stronger signal in WT compared to the *gaut1-1/gaut1-1*. These results suggested that the doublet bands represented either a non-HG moiety or a very highly modified HG since they were resistant to EPG treatment. We suspected RG-II monomer as the likely candidate for these bands due to (1) the alignment of bands with the presumed RG-II monomer in the sodium carbonate fractions, and (2) the likely monomerization of the borate-diester dimer of RG-II in the 4M KOH extraction due to the strong base. The downregulation of the doublet bands in *gaut1-1/gaut1-1* was most prominent in the final wall extract i.e. the 4M KOH post-chlorite fraction, where the bands were nearly absent in the *gaut1-1/gaut1-1* sample compared to WT. As mentioned above, the 4M KOH post-chlorite was the only wall fraction where GalA content was significantly reduced in the *gaut1-1/gaut1-1* compared to WT (Table 2). The absence of the band in the 4M KOH post-chlorite fraction of the *gaut1-1/gaut1-1* mutant further suggested that the doublet bands were either not synthesized or not held in the wall in the same manner when the HG synthesized by GAUT1 is not produced.

To determine the identity of the EPG-resistant polymer band, aliquots of the EPG-digested 4M KOH fraction (chosen because it provided the most material) were separated by anion exchange chromatography over a diethylaminoethyl (DEAE) column followed by two rounds of size exclusion chromatography over a Superdex Peptide column. Aliquots of fractions collected from each column were tested for the presence of the bands by HP-PAGE (Figure S11; also see Materials and Methods for details). The band-containing fractions were pooled and passed through a solid phase extraction graphite column and the band material then eluted with 50% acetonitrile. NMR spectroscopy of this purified material confirmed its identity as RG-II, based on the similarities between the ^13^C-HSQC spectrum of purified band material and the ^13^C-HSQC of a known RG-II sample from red wine (Figure S12). Characteristic signals from multiple side-chain residues unique to RG-II (Herve du Penhoat et al., 1999; Glushka et al., 2003; Rodriguez-Carvajal et al., 2003) were clearly observed. For example, the methyl and methylene region of the HSQC spectrum (Figure S12B) contains signals for multiple rhamnoses, fucoses, aceric acid, Dha (3-deoxy-D-*lyxo*-2-heptulosaric acid) and Kdo (2-keto-3-deoxy-D-*manno*-octulosonic acid). Similarly, the anomeric (H_1_, C_1_) region (Figure S12C) shows characteristic signals for α-L-galactopyranose, α-L-2-*O*-Me-fucose, D-apiose and L-arabinofuranoses, all sugars that are unique to RG-II. Some differences in chemical shifts between the spectra were also observed. However, this is not uncommon for different preparations and different sources of RG-II due to structural heterogeneity and sample conditions. The red wine sample is a borate cross-linked dimer that is acetylated, thus it has greater molecular weight and heterogeneity, while the Arabidopsis EPG-resistant target band is more consistent with an RG-II monomer based on the lineshape. Taken together, these results suggested that the HG synthesized by GAUT1 serves as the backbone for a subfraction of cell wall RG-II, and that this HG-RG-II-HG heteropolymer subpopulation that is tightly bound in the wall was downregulated in the *gaut1-1/gaut1-1* mutant.

### dSTORM reveals reduced formation of cell wall HG nanofilaments in *gaut1-1/gaut1-1* callus and seedling

Taken together the results indicated that a specific HG-associated cell wall polymer synthesized by GAUT1 was essential for proper cell expansion/elongation in Arabidopsis. Pectic HG nanofilaments have been identified previously as components involved in cell expansion control in Arabidopsis epidermal cells (Haas et al., 2020a). Specifically, super-resolution microscopy combined with immunolabelling revealed that highly methylesterified and low-methylesterified HG, coordinated by calcium ions and detected respectively by LM20 and 2F4 antibodies, can assemble into short nanofilaments within the anticlinal walls of Arabidopsis cotyledons. To determine if HG nanofilaments were affected in the *gaut1-1/gaut1-1* mutant, seedlings and calli from WT and *gaut1-1/gaut1-1* mutant were analyzed by super-resolution three-dimensional direct stochastic optical reconstruction microscopy (3D-dSTORM) (Huang et al., 2008; Haas et al., 2020a) combined with immunolabelling using LM20 and 2F4 antibodies (Figure 9). Given the severely impaired growth of homozygous *gaut1-1/gaut1-1* seedlings, the *gaut1-1/gaut1-1* mutant seedlings were compared to WT Col-0 seedlings at equivalent developmental stages.

**Figure 9.**
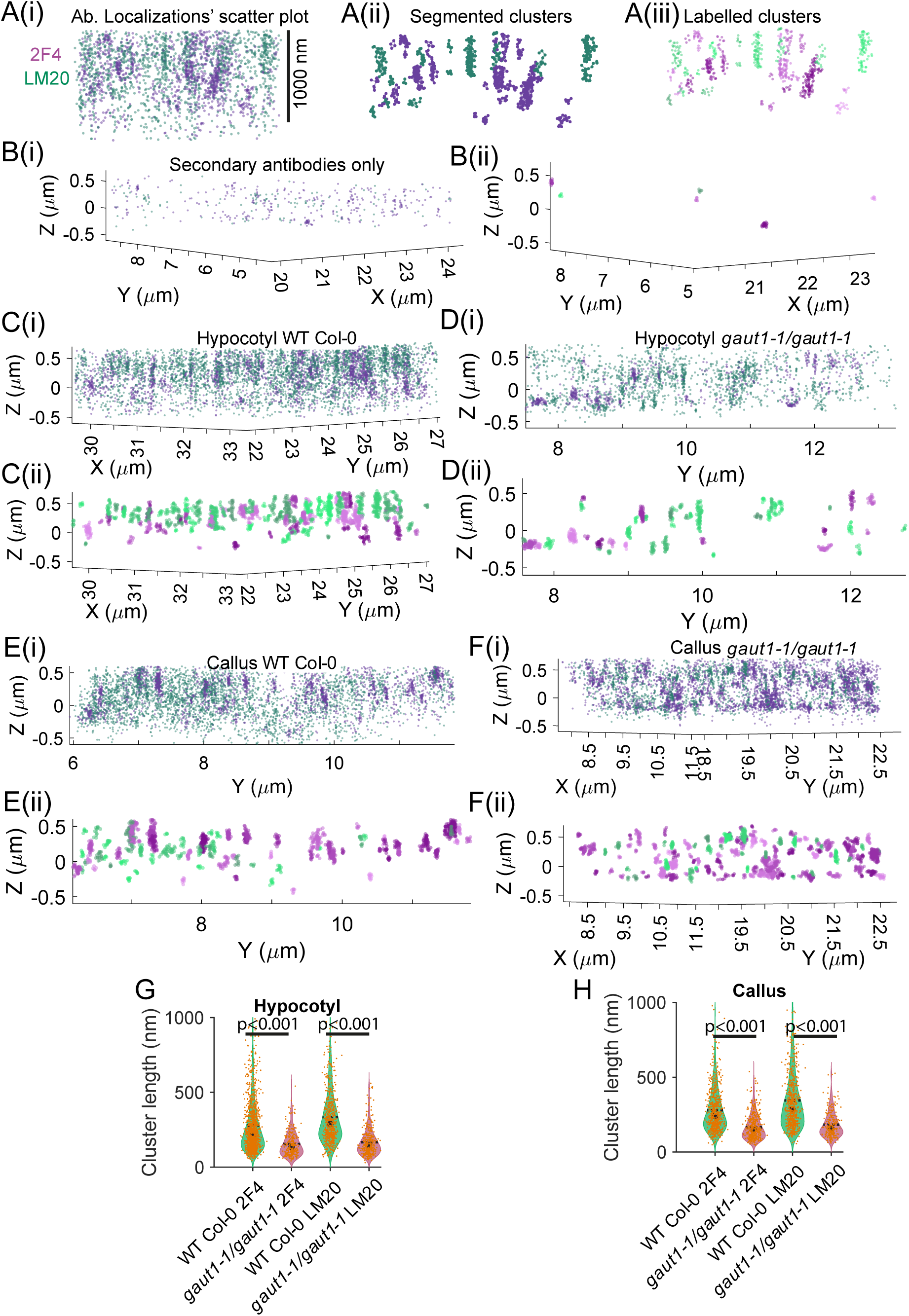
dSTORM analysis revealed significantly reduced pectin nanofilaments in *gaut1-1/gaut1-1* mutant cell walls in comparison to WT (Col-0). (A(i)) Side-on 3D view of a representative dSTORM scatter plot of the hypocotyl anticlinal wall in the Col-0 background, immunolabeled for methylated homogalacturonan using the LM20 antibody (Ab) (green), showing a discontinuous fibrous pattern. **(A(ii))** The data from (A(ii)) is segmented using 3D Delaunay triangulation resulting in disconnected graphs i.e. nodes connected by edges, where all edges longer than a threshold were removed, see the Materials and Methods. **(A(iii))** Labelled disconnected clusters, only nodes are displayed. **(B(i))** dSTORM scatter plot of a control sample tagged with secondary antibodies only. **(B(ii))** Clusterization of the data from **(B(i))**. **(C(i))** Side-on 3D view of a representative dSTORM scatter plot of the hypocotyl anticlinal wall in the WT Col-0 background, immunolabeled for methylated (green) and demethylated (purple) homogalacturonan using LM20 and 2F4 antibodies, respectively, highlighting the visible fibrous pattern. **(C(ii))** Segmented clusters from **(C(i))**. **(D(i))** Side-on 3D view of a representative dSTORM scatter plot of the hypocotyl anticlinal wall in the *gaut1-1/gaut1-1* background, immunolabeled for methylated (green) and demethylated (purple) homogalacturonan using LM20 and 2F4 antibodies, respectively. **(D(ii))** Segmented clusters from **(D(i))**. **(E(i))** Side-on 3D view of a representative dSTORM scatter plot of the callus cell wall in the WT Col-0 background, immunolabeled for methylated (green) and demethylated (purple) homogalacturonan using LM20 and 2F4 antibodies, respectively. **(E(ii))** Segmented clusters from **(E(i))**. **(F(i))** Side-on 3D view of a representative dSTORM scatter plot of the callus anticlinal wall in the *gaut1-1/gaut1-1* background, immunolabeled for methylated (green) and demethylated (purple) homogalacturonan using LM20 and 2F4 antibodies, respectively. **(F(ii))** Segmented clusters from **(F(i))**. **(G, H)** Violin plots showing segmented LM20 and 2F4 cluster lengths in Col-0 (green) and *gaut1-1/gaut1-1* (bordeaux) backgrounds, observed in the calli and hypocotyls, respectively. Orange dots represent individual cluster data points. *P*-values were obtained using the Kruskal–Wallis test with Bonferroni correction. In B(ii), C(ii), D(ii) the segmented LM20 clusters are labelled using light-to-dark green colormap and 2F4 light-to-dark purple colormap. Number of (2F4, LM20) clusters analysed in different conditions: (2361, 1225) in WT Col-0 seedlings, (614, 1010) in *gaut1-1/gaut1-1* seedlings, (1391, 1611) in WT Col-0 calli, and (1236, 780) in *gaut1-1/gaut1-1* calli. The clusters with a minimum 10 detections after clustering were retained for the final analyses.

For quantitative analysis of nanofilament occurrence, the MATLAB-based Grafeo software was used (Haas and Peaucelle, 2021) (Figure 9A). The dSTORM data consist of 3D coordinate lists of localized fluorophores conjugated to the antibodies, typically visualized as scatter plots of 3D point clouds (Figure 9A(i)). To segment these localizations into discrete clusters, Delaunay triangulation-based thresholding was applied, wherein 3D points (nodes) were connected by edges representing Delaunay triangles. Edges exceeding a threshold length (typically ∼50 nm; see Materials and Methods) were removed, resulting in a discontinuous tessellation that isolates clusters (Figure 9A(ii)). To enhance visualization, clusters were color-coded: LM20-labelled clusters in shades of green, and 2F4-labelled clusters in purple (Figure 9A(iii)).

Control labelling with only secondary antibodies, in the absence of primary antibodies, produced dispersed localizations (Figure 9B(i)) with very few small, segmented clusters (Fig. 9B(ii)), confirming labelling specificity. In WT seedlings, dSTORM imaging revealed dense bands resembling pectin nanofilaments (Figure 9C(i)), consistent with those previously reported in pavement cells (Haas et al., 2020a). Cluster segmentation effectively excluded dispersed localizations, highlighting elongated, rod-like pectin nanofilaments (Figure 9C(ii)). In contrast, *gaut1-1/gaut1-1* seedlings exhibited visibly shorter and more isotropic pectin clusters (Figure 9D). In WT calli, we also observed elongated HG clusters (Figure 9E), which were largely absent in *gaut1-1/gaut1-1* calli (Figure 9F). Finally, violin density plots comparing segmented cluster lengths across conditions confirmed a significant reduction in nanofilament length in *gaut1-1/gaut1-1* backgrounds relative to WT Col-0 (Figures 9G, H), supporting the hypothesis that GAUT1 synthesizes HG glycans essential for nanofilament formation.

## DISCUSSION

The biochemical characteristics of Arabidopsis GAUT1, the first identified HG:GalAT, have been extensively explored, yet the exact nature of the GAUT1-synthesized-HG-containing polymeric product and its biological and biochemical function(s) *in vivo* have remained elusive. Here characterization of an Arabidopsis homozygous *gaut1-1/gaut1-1* T-DNA insertion mutant revealed that the HG synthesized by GAUT1 is required for both vegetative and reproductive stages of plant development. The absence of GAUT1 results in severely stunted seedlings of *gaut1-1/gaut1-1* homozygous mutants (Figures 1 and 2), whose frequency of occurrence is substantially reduced (Table 1) due to defective growth of pollen tubes from heterozygous plants and to defective embryo/seed development in homozygous individuals (Figure 4). At the cellular level, GAUT1 product is critical for proper cell expansion/elongation (Figures 2 and 3), but does not appear essential for cell division, differentiation, or for tissue organization (Figure 3). These results suggest that there are different and unique HG moieties in the plant cell wall and that these have unique functions/roles. The results also indicate that GAUT1 synthesizes HG that is a part of a wall polymer and architectural component required for both diffusive cell expansion/elongation and for tip growth. GAUT1 function *in planta* appears to be unique and non-redundant with the function(s) of other GAUTs as it was not compensated for by any other GAUT since no mature homozygous *gaut1-1/gaut1-1* plants were isolated despite upregulation of the transcript expression of other GAUTs in the *gaut1-1/gaut1-1* seedlings. Lund et al. (2020) also reported that the Arabidopsis triple homozygous mutant of GAUT5, GAUT6, and GAUT7, GAUTs which interchangeably serve as the membrane anchors of GAUT1, was not obtainable. These data confirm the proposition that different GAUTs have unique functions in plants and that GAUT1 has a singular unique function, at least in Arabidopsis, since no mature homozygous *gaut1-1/gaut1-1* mutant plants could be recovered.

Considering the severe penalty on vegetative growth due to the absence of GAUT1, it is surprising that *gaut1-1/gaut1-1* callus and suspension cultures could be generated and thrive. The report by Frankevich et al. (2025) on the generation and analysis of suspension cell cultures of a CRISPR-knocked out (KO) mutant of Arabidopsis GAUT1, described some similar characteristics to the *gaut1-1/gaut1-1* cultures studied here. Specifically, Frankevich et al. (2025) reported darker coloration, larger cell aggregates, slightly slower growth rates, and more extracellular material being exuded in GAUT1 CRISPR-KO mutant suspension cell cultures compared to control. The GAUT1 CRISPR-KO calli were also reported to have a loose structure, which the authors suggested was due to faulty cell-to-cell adhesion. However, that observation does not agree with the larger cell aggregates seen in the GAUT1 CRISPR-KO suspension culture, and is contrary to the results obtained here in which the *gaut1-1/gaut1-1* calli were less friable than WT and based on ruthenium red staining of *gaut1*-/- seedling hypocotyls showing no defective cell-to-cell adhesion due to the absence of GAUT1. Moreover, Frankevich et al. (2025) reported no difference in pectin content between the GAUT1 CRISPR-KO suspension cells compared to control, which is also contrary to our findings here in which a modest but significantly reduced GalA content was observed in the *gaut1-1/gaut1-1* suspension cultured cell walls (Figure 6A). This discrepancy could be due to the different culture conditions used to grow the suspension cultures as well as to the different methods used to measure pectin content in the Frankevich et al. (2025) study versus this work. Intriguingly, we report here that the large amount of extracellular polymer extruded by *gaut1-1/gaut1-1* culture actually contains elevated amounts of GalA, presumably HG (Figure S6). One plausible reason for the increased HG in the *gaut1-1/gaut1-1* mutant suspension culture medium reported here is that the loss of GAUT1-synthesized HG results in a looser cell wall that retains less of the synthesized wall polysaccharides (hence its extrusion into apoplast in the case of callus and into the liquid medium in the case of suspension cultured cells). The results of the glycome profiling experiments (Figure 7), which show looser wall association of xyloglucan and galactan epitopes, support this hypothesis. It is also possible that HG oligosaccharides not held in the wall could serve as signals to the cells to produce more pectin and other wall polysaccharides (Voxeur and Hofte, 2016).

GAUT1 is a major contributor of HG:GalAT activity in Arabidopsis suspension cultures, responsible for about one fourth of total HG:GalAT activity. Although GAUT1 is a highly enzymatically active GAUT that synthesizes high molecular weight HG chains *in vitro* (Amos et al., 2018), the *gaut1-1/gaut1-1* mutant enzyme activity and cell wall data clearly indicate that other HG:GalATs, presumably other GAUTs, must also be expressed and present in Arabidopsis suspension culture cells and synthesize HG. Indeed, quantitative PCR transcript expression analysis of the 15 Arabidopsis GAUTs (Figure S7) showed that all of them, with the exception of GAUT2 and GAUT12, are expressed at various levels in Arabidopsis suspension cultures. As mentioned above, however, relatively high expression levels (comparable to *GAUT1* transcript levels) of *GAUTs 4, 8, 9* and *15* in WT culture were observed, suggesting that the remaining three quarters of total HG:GalAT activity is likely largely attributable to these particular *GAUTs*. In fact, GAUT4 has previously been shown to synthesize the bulk of HG in poplar, switchgrass and rice, since downregulation of its transcript expression by 49-78% resulted in 74-91% reduction in the total wall GalA content and an increase in cell size (Biswal et al., 2018b). In contrast, the abolished transcript expression of *GAUT1* in the *gaut1-1/gaut1-1* mutant suspension culture resulted in about 25% reduction in the total HG:GalAT activity and total GalA content and a decrease in cell size. These results suggest that GAUT1-synthesized HG exerts its function due to a specific HG polymer structure that is different than GAUT4-synthesized HG.

Analysis of sequentially fractionated cell walls of *gaut1-1/gaut1-1* and control suspension culture lines revealed a significant reduction in GalA in the *gaut1-1/gaut1-1* mutant 4M KOH post-chlorite cell wall fraction, which was surprising since this final wall fraction is generally expected to contain cell wall polymers held tightly in the wall, possibly via crosslinks with lignin. The large and significant reduction of GalA in the 4M KOH post-chlorite fraction of the *gaut1-1/gaut1-1* mutant compared to WT suggests that GAUT1 synthesizes an HG that is present in a polymer that is tightly bound in the wall requiring harsh solvents for its release. It is possible that GAUT1-synthesized HG is part of the so-called insoluble pectin previously reported by Selvendran (1985). The slight reduction of all other sugar residues, with the exception of Ara, as revealed by sugar composition analysis of the 4M KOH post-chlorite fraction of the *gaut1-1/gaut1-1* mutant may indicate an association of GAUT1-synthesized HG with other glycans in the hemicellulose-cellulose network, such as xyloglucans, xylans, mannans, or cellulose, or with lignin. Such associations would explain the tight binding of GAUT1-synthesized HG to the cell walls, thus necessitating the use of both acidified sodium chlorite treatment to break lignin/phenolic connections and strong base treatment to break the ester- and/or hydrogen bond connections (Selvendran, 1985; Broxterman et al., 2018; Shakhmatov and Makarova, 2022). This contrasts with the above-mentioned GAUT4, which synthesizes the bulk of HG (Biswal et al., 2018b), and with GAUT12, which synthesizes HG required for xylan synthesis and/or deposition (Hao et al., 2014; Biswal et al., 2015; Biswal et al., 2018a). With both GAUT4 and GAUT12, their downregulation causes GalA content reduction in virtually all sequentially extracted wall fractions (Biswal et al., 2015; Biswal et al., 2018b), rather than a reduction in only a specific fraction, as occurs with the Arabidopsis *gaut1-1/gaut1-1* mutant. These results further support the hypothesis that different GAUTs synthesize different HG glycans, and that GAUT1 synthesizes a unique HG tightly held in the wall, presumably due to association with other wall glycans or constituents.

The EPG treatment of the wall fractions sequentially extracted from WT and *gaut1-1/gaut1-1* suspension culture cells followed by visualization of the hydrolysates on HP-PAGE identified the connection of GAUT1-synthesized HG with RG-II. Classically, the bulk of RG-II is isolated from cell walls by treating total AIR or the pectin-rich ammonium oxalate fraction with EPG, followed by size exclusion chromatography to separate RG-II from RG-I and HG oligomers (Darvill et al., 1978; Barnes et al., 2021; Hays et al., 2025). Indeed, our results confirm that the bulk of RG-II appears to be extracted in the ammonium oxalate fraction and to a lesser extent in the sodium carbonate fraction (Figure 8B). However, we found here, unexpectedly, that RG-II also exists in wall fractions extracted by the harshest solvents, i.e. 4M KOH, acidified sodium chlorite, and 4M KOH post-chlorite. These results support the premise that all the major classes of pectins (HG, RG-I, and RG-II) are present in the walls in diverse polymer populations, including polymers that are extracted in water soluble fractions and polymers that are extracted only/largely in the most tightly bound cell wall fractions (Mohnen et al., 2024). The data also suggest that GAUT1, together with the previously reported GAUT4 (Biswal et al., 2018b), synthesize HG backbone onto which the RG-II side chains are added by a yet-to-be-elucidated biosynthetic process. For example, it is not known whether the RG-II side chains are added one residue at a time directly onto the HG backbone, or are first synthesized on a separate primer and then transferred as whole side chains onto the HG backbone. Furthermore, the actual structure of the whole heteropolymer, prior to EPG digestion, remains elusive and requires further investigation. It is likely that the HG-RG-II-HG structure is connected to as yet unidentified other glycans and/or phenolic compounds that are removed together with HG upon EPG treatment. Elucidation of the full polymeric structure in which the GAUT1-synthesized HG resides is critical for determination of how this particular HG- and RG-II-containing polymer organizes into HG nanofilaments associated with cell expansion.

The results reported here provide multiple lines of evidence indicating that the loss of GAUT1 leads to altered cell wall architecture with both a loss of HG nanofilaments and a looser cell wall architecture that does not hold wall material properly in the wall, resulting in earlier release of wall polymers during sequential cell wall extractions. The evidence includes a greater amount of fibrous material on the surface of *gaut1-1/gaut1-1* calli and a greater amount of ECM released into the *gaut1-1/gaut1-1* suspension culture media. Notably, the earlier detection, compared to WT, of xyloglucan epitopes in the *gaut1-1/gaut1-1* ammonium oxalate, sodium carbonate, and 1M KOH fractions as revealed by glycome profiling analysis was unexpected. It is plausible that GAUT1-synthesized HG might be involved in crosslinking xyloglucans in the wall. Polysaccharide complexes containing both pectins and xyloglucans have previously been described (Chambat et al., 1984; Cornuault et al., 2018). Alternatively, it is possible that the loss of the GAUT1-synthesized HG leads to such an extensive disruption of the overall wall structure such that xyloglucans can no longer be properly fixed into the wall structure.

It has been previously shown by super resolution microscopy that at least some HG in growing cells, for example in Arabidopsis cotyledon pavement cells, resides in HG nanofilaments that function in controlling cell expansion (Haas et al., 2020a). However, the mechanisms underlying the formation, composition, and stability of these HG nanofilaments remain unclear, including whether additional molecular components are required. Interestingly, HG nanofilaments were not observed in the anticlinal walls of Arabidopsis pavement cells when immunolabelled with the JIM7 antibody, which recognizes partially methylesterified HG (Haas et al., 2021). This suggests that the pattern of HG methylesterification, possibly shaped by the mode of pectin methylesterase (PME) activity (processive vs. non-processive), may play a critical role in nanofilament formation. Additional insights came from dSTORM imaging combined with immunolabelling using the LM19 antibody, which recognizes low-methylesterified HG, in Arabidopsis pollen tubes (Moussu et al., 2023). In these cells, LM19-labelled HG assembled into a crisscrossed fibrous network, whose formation depended on the presence of the LEUCINE-RICH REPEAT EXTENSIN 8 - RAPID ALKALINIZATION FACTOR 4 (LRX8-RALF4) complex. This finding suggests that proteinaceous components are crucial for the structural stability of HG nanofilaments. Similarly, a basal pectic fibrous pattern was observed in the unicellular charophyte *Penium margaritaceum* through both immunolabelling with the JIM5 antibody, which recognizes low-methylesterified HG, and electron microscopy (Palacio-Lopez et al., 2020).

Evidence for fibrillar pectin structures dates back to the 1950s. Electron microscopy studies reported fibrillar pectin in plants, including observations of axially aligned pectin crystallites within fresh and partially dried collenchyma cell walls of *Petasites vulgaris* petioles, as revealed by distinct meridional diffraction patterns in X-ray diffraction analyses, differing from those of cellulose (Roelofsen and Kreger, 1951). The scarcity of data on fibrillar pectin structures suggests that such assemblies may be tightly integrated within the cell wall matrix and thus potentially masked or degraded during conventional extraction protocols. Alternatively, pectin nanofilaments may be inherently unstable and therefore disrupted during isolation. Here we asked whether the HG synthesized by GAUT1 might be associated with HG nanofilaments. We show that the formation of HG nanofilaments is greatly reduced both in the walls of *gaut1-1/gaut1-1* seedling hypocotyl epidermal cells and in the walls of *gaut1-1/gaut1-1* callus cells. From these results, we hypothesize that GAUT1 synthesizes HG that is a component of and required to organize pectins into nanofilaments in the wall, and that the GAUT1-synthesized HG-containing polymer in the nanofilaments has both HG and RG-II domains. When the GAUT1-synthesized HG is not made, both HG nanofilament length and cell expansion are greatly reduced. Taken together, the results lead us to propose that at least in higher plants GAUT1 plays a fundamental role in cell expansion by synthesizing HG that is a component of nanofilaments that are required for proper cell expansion/elongation (Figure 10). Further critical questions arise from this proposition. For example, what is the precise role of RG-II in pectin nanofilament structure and function? How long is the HG in the nanofilament and how is this length regulated? How is pectin organized in the nanofilament? Are non-pectic polymers constituents of the nanofilaments? Do the nanofilaments have a protein/peptide component? Further research is required to provide answers to these questions.

**Figure 10.**
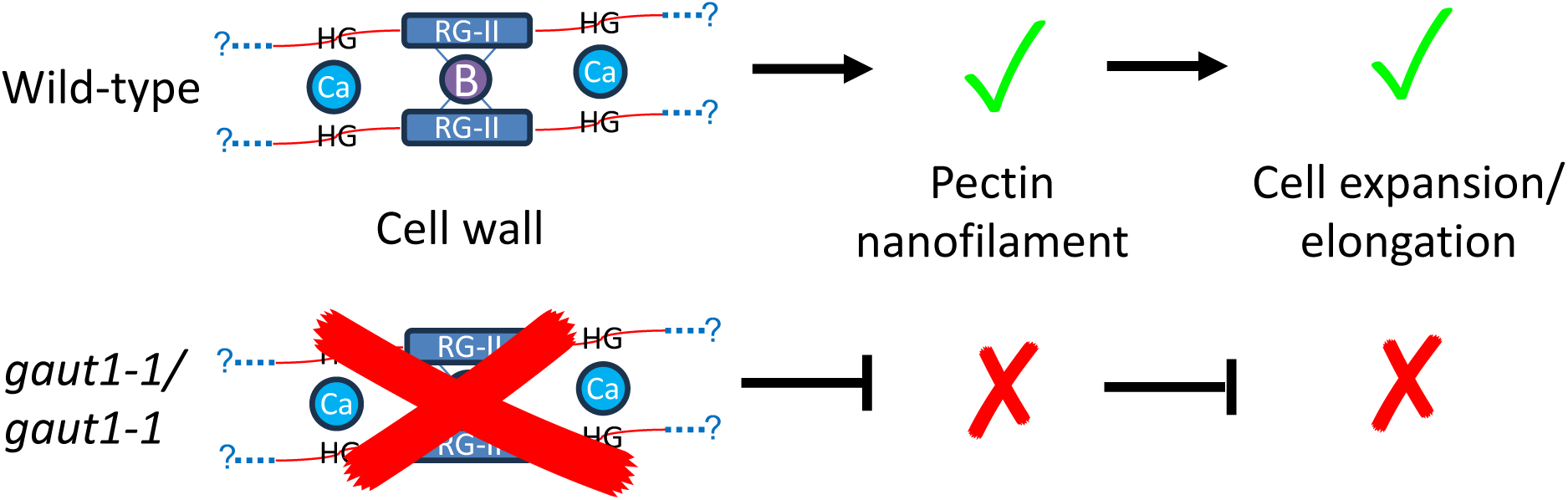
Hypothetical model of Arabidopsis GAUT1-synthesized HG as a major component of HG nanofilaments and associated reduced cell expansion in *gaut1-1/gaut1-1* mutants. GAUT1 synthesizes HG present in HG nanofilaments that are tightly held in the wall and associated with cell expansion. The HG synthesized by GAUT1 serves as the backbone of a species of RG-II present in nanofilaments, and might be further connected to unknown, to-be-determine polymer(s) (blue dotted lines with question marks). In the *gaut1-1/gaut1-1* mutant, lack of GAUT1-synthesized HG results in a lack of, or drastic reduction in, HG nanofilaments and a lack of HG nanofilaments-associated cell expansion.

Finally, the results of this molecular genetic characterization of the *gaut1* mutant provide conclusive data that different GAUTs synthesize HG that resides in polymers that reside in different fractions and structures in the plant cell wall. Thus, current cell wall models that present pectin as only HG, RG-I, or RG-II polysaccharides are incomplete. As we gain an understanding of the pectic domains that reside in specific polymers, more accurate polymer-specific models can be made (akin to models of the specific proteoglycans and glycosaminoglycans that exist in animal extracellular matrices and carry out specific functions, exemplified by aggrecan (Hayes and Melrose, 2020). At present, we name the GAUT1-synthesized HG-containing polymer as GAUT1-HG-RG-II polymer. This name will likely need modification as we understand more about the full structure of the polymer and whether it has non-pectin component(s).

## MATERIALS & METHODS

### Materials

All chemicals, unless otherwise noted, were from Sigma-Aldrich Corp. (St. Louis, MO). Quantitative real-time PCR reagents were from Bio-Rad (Hercules, CA). Uridine diphosphate α-D-[^14^C] galacturonic acid (UDP-D-[^14^C]Gal*p*A) was synthesized as previously described (Atmodjo et al., 2011) from UDP-D-[^14^C]Glc*p*A (6.67 GBq/mmol) purchased from PerkinElmer. Uridine diphosphate α-D-galacturonic acid (UDP-D-Gal*p*A) was from Carbosource Services (Athens, GA). *Arabidopsis thaliana* var. Columbia T-DNA insertion mutant line GK-470H07-019028 was obtained from the Nottingham Arabidopsis Stock Centre (NASC, http://arabidopsis.info), while the lines SAIL_1162_A03, WiscDsLoxHs190_06B, SALK_063880, and SALK_013652 were obtained from the Arabidopsis Biological Resource Center (ABRC, https://abrc.osu.edu/). The plant cell wall glycan-directed monoclonal antibodies used in this study were those described in Pattathil et al. (2010) and are obtainable from CarboSource Services (Athens, GA). Detailed information about the epitopes recognized by these antibodies is summarized in a recent review (Thorne et al., 2024).

### Plant growth conditions

For *in vitro* seedling observation, sterilized wild type and mutant seeds were stratified for 2-4 days and germinated on agar plates containing half-strength Murashige and Skoog basal medium (pH 5.8) and 5.5 g/L plant agar (Research Products International Corp., Mount Prospect, IL), and kept in a growth chamber with 60% constant relative humidity and a photoperiod cycle of 14/10 hr light/dark (150 μmol photons m^-2^ s−_1_ light) at 19°C/15°C. For plant observation, stratified seeds were sown directly on pre-moistened soil supplemented with Osmocote (Outdoor and Indoor; The Scotts Company, Marysville, OH) following the manufacturer’s instruction, and grown to maturity in a growth chamber under the same conditions as above except for a 16/8 hr light/dark cycle.

### Genomic DNA, transcript and protein analyses

For genotyping, genomic DNA samples were prepared from Arabidopsis plants using the REDExtract-N-Amp Plant PCR kit (Sigma-Aldrich, St. Louis, MO), and from calli using PowerPlant DNA isolation kit (Mo Bio Lab. Inc., Carlsbad, CA). The presence of the T-DNA insertion within the Arabidopsis *GAUT1* (*At3g61130*) gene was determined using primers specific to the genomic regions flanking the T-DNA insertion sites and a T-DNA left border primer (Table S2). For transcript analysis, RNA was extracted from plant samples using the RNeasy Plant Mini Kit (Qiagen, Germantown, MD) and assessed using a Nanodrop spectrophotometer (Thermo Fisher Sci., Waltham, MA). cDNA was synthesized from 1 μg extracted RNA using Bio-Rad iScript Reverse Transcription Supermix for RT-PCR, and the real time quantitative PCR carried out using Bio-Rad iQ SYBR Green Supermix with primers as listed in Table S2. For protein analyses, microsome preparations from the suspension cultured cells, Western blots, and HG:GalAT activity assays were carried out as previously described (Biswal et al., 2018b).

### Cloning and plant transformation to generate complementation line

*GAUT1* coding sequence (CDS) was amplified from WT leaf cDNA using primers G1-CL-1F and G1-CL-1R (Table S2) that also introduced SmaI and SacI restriction sites at the 5’-and 3’-end of the CDS, respectively. Following cloning into pGEM-T vector (Promega) and verification by DNA sequencing, the *GAUT1* CDS was subcloned into plant expression plasmid pBI101-*ProGAUT1* (Atmodjo et al., 2011), replacing the original GUS gene downstream of GAUT1 promoter sequence. *Agrobacterium tumefaciens* (strain GV3101::pMP90) harboring the resulting construct of GAUT1-CDS driven by GAUT1 promoter (pBI101-*ProGAUT1:GAUT1*), was used to transform Arabidopsis heterozygous *GAUT1/gaut1-1* plants using the floral dip method (Clough and Bent, 1998). Complemented homozygous mutants were identified by genotyping the T1 population for both the homozygous *gaut1-1/gaut1-1* mutant background and the presence of the introduced *GAUT1* CDS.

### Microscopy

Seedling growth was observed using a dissecting scope (Olympus SZH) and pictures taken using a Nikon Ds-Ri1 digital camera. For scanning electron microscopy, WT or *GAUT1*/*gaut1-1* and *gaut1-1/gaut1-1* seedlings were fixed in 2.5% (w/v) glutaraldehyde (Electron Microcopy Sciences, Hatfield, PA) in 25 mM sodium phosphate buffer pH 7.1 and 0.02% (v/v) Tween-20 for 2 hr at room temperature (RT). Seedlings were washed three times with ddH_2_O for 20 min each at RT and then subjected to an ethanol gradient incubation series (25, 50, 75, 95, 100, and 100% (v/v), 20-30 min each at RT). After critical point drying, seedling specimens were mounted onto specimen stubs covered with adhesive graphite tabs, coated with gold palladium alloy particles using a Leica EM ACE200 (Leica Microsystems Inc., Buffalo Grove, IL) vacuum coater, and observed using JEOL JSM-6010LV scanning electron microscope with InTouchScope software (JEOL USA Inc., Peabody, MA). Size measurements from captured images were done using ImageJ (https://imagej.nih.gov/ij/).

For light microscopy of seedling hypocotyl cross sections, specimens were prepared essentially as described (Biswal et al., 2015; Majda, 2019). Seven-day-old WT and *gaut1-1/gaut1-1* seedlings were incubated in fixing solution (4% (w/v) paraformaldehyde, 0.5% (w/v) glutaraldehyde (Electron Microscopy Sciences, Hatfield, PA), 0.02% (v/v) Triton X-100 in 25 mM sodium phosphate buffer pH 7.1) for several hours at RT followed by overnight at 4°C, then washed for 30 min each twice with sodium phosphate buffer (25 mM, pH 7.1) and twice with ddH_2_O. Specimens were subsequently embedded in low melting point agarose, passed through a graded ethanol series, and embedded in LR White resin (Ted Pella) as described (Majda, 2019). Sectioning of the specimens into 250 nm sections was carried out by the Georgia Electron Microscopy facility at the University of Georgia (Athens, GA). Light microscopy observation and documentation were done using a Nikon Eclipse 80i microscope equipped with a Nikon DS-Ri2 camera (Nikon, Melville, NY).

For size measurement of the suspension cultured cells, clumps of cells from 7-day-old cultures were macerated as previously described (Biswal et al., 2015) with incubation for 3 hours at 100°C in 1 mL solution of 50% (v/v) glacial acetic acid and 3% (v/v) hydrogen peroxide. Light microscopic observation of the individual cells was carried out as described above and the cell size was measured from photomicrographs using ImageJ (https://imagej.nih.gov/ij/).

### Pollen viability, germination and tube growth

Assessments of pollen viability, germination, and tube growth were carried out following protocols as described (Lund et al., 2020). Pollen tube length was determined from photomicrographs using ImageJ (https://imagej.nih.gov/ij/).

### Generation, maintenance, and collection of suspension cultured cell lines

Segregating seeds from a *GAUT1/gaut1-1* heterozygous plant were sterilized, cold-treated at 4°C for 2 days, and sown directly on suspension culture medium (Guillaumie et al., 2003) solidified with 5.5 g/L plant agar. Plates were kept in the dark at RT. Seeds were allowed to germinate and form calli, which were subcultured onto fresh medium plates approximately every 3-4 weeks. Calli were genotyped by PCR and representative WT, heterozygous, and homozygous lines were isolated. Three-week-old calli were used to inoculate suspension culture medium (without plant agar) in flasks with shaking (100 rpm) to generate the suspension cultures. The cultures were regularly maintained by subculturing 6 mL packed cells into 100 mL fresh medium every 10 days. Cultures for growth experiments and other analyses were prepared the same way, except that 5 mL packed cells were used for every 100 mL medium. The cultures were harvested at day 7 (unless otherwise noted) by filtering through a nylon mesh (100 μm). The retained cells were rinsed twice with deionized water, flash frozen in liquid nitrogen, and kept at -80°C until use. The flow-through was passed through a filter paper (Whatman no. 1) to obtain cleared culture medium containing extracellular material (ECM), which was subsequently dialyzed, lyophilized, and kept at RT until use.

### Cell wall preparation and analyses

Frozen suspension cultured cells (∼5 g) were ground to a fine powder in liquid N_2_ and extracted sequentially in 50 mL volume of 80% (v/v) ethanol, 96% (v/v) ethanol, 1:1 (v/v) chloroform:methanol, and 100% acetone, each for 8 hr to overnight at RT with constant end-to-end rotation. The resulting alcohol insoluble residue (AIR) samples were air dried in a fume hood for 1 to 2 days. AIR samples (300 mg) were then treated with 285 units of porcine pancreas α-amylase in 30 mL of 100 mM ammonium formate buffer, pH 6.0 with 0.01% (w/v) thimerosal for 48 hr at RT with constant end-to-end rotation, washed three times with ddH_2_O and twice with 100% acetone, and air dried in a fume hood to yield destarched AIR (dsAIR) samples. Cell wall fractionation by sequential extractions using increasingly harsh solvents was carried out on 200 mg dsAIR following procedures as described (Pattathil et al., 2012).

Glycosyl residue composition was determined from dsAIR (2 mg) and wall fractions (0.5 to 1 mg) using gas chromatography – mass spectrometry (GC-MS) of trimethylsilyl (TMS) derivatives essentially as described (Biswal et al., 2015) with 18 hr and 16 hr methanolysis for dsAIR and wall fractions, respectively. The same method was also used to determine cellulose content of dsAIR following Saeman hydrolysis (Saeman et al., 1945). Glycosyl linkage analysis was performed as previously described (York et al., 1985; Black et al., 2023). For methyl and acetyl content determination, 1 mg dsAIR, ammonium oxalate fraction, or sodium carbonate fraction samples were saponified in 600 μl 0.5 N NaOH for 1 hr at 4°C, neutralized by addition of 300 μl 1N HCl, and assayed using BioVision (Milpitas, CA) methanol assay and Megazyme (Wicklow, Ireland) acetic acid assay kits, respectively. Boron and calcium contents were determined from 20 mg dsAIR or 2 to 3 mg of the ammonium oxalate and sodium carbonate fractions by Inductively-Coupled Plasma – Optical Emission Spectroscopy (ICP-OES) at the Center for Applied Isotope Studies, University of Georgia, Athens. Glycome profiling of cell wall extracts was carried out as previously described (Pattathil et al., 2012).

### HP-PAGE, chromatography, and NMR

Alcian blue-silver nitrate-stained HP-PAGE was carried out as previously described (Amos et al., 2018). For digestion with endopolygalacturonase (EPG), the wall fractions were treated in a reaction containing 10 mg/mL wall fractions and 0.005 mg/mL EPG (EPG-I from *Aspergillus niger*, EC 3.2.1.15) in 33.3 mM sodium acetate, pH 5.0 that was incubated at 40°C overnight.

The WT EPG-digested 4M KOH wall fraction was further separated on a gravity anion exchange column of diethylaminoethyl (DEAE) Sephacel (Cytiva, Wilmington, DE), eluted with a step gradient of ammonium formate (100 mM, 200 mM, 300 mM, 400 mM, 500 mM, and 2 M). The fractions obtained were dialyzed (1000 Da MWCO) against water and lyophilized to dryness. A small aliquot of each fraction was electrophoresed on a high percentage polyacrylamide gel (HP-PAGE) to monitor the presence of the EPG-resistant target bands. Fractions number 3 to 6 from the DEAE column were found to contain the target band (Figure S11A) and these were combined and subsequently separated by size exclusion chromatography (SEC) over a Superdex Peptide 10/300 GL column (Cytiva, Wilmington, DE) connected to an HPLC (Shimadzu) equipped with refractive index and UV (220 and 280 nm) detectors, using 50 mM ammonium formate as the mobile phase and a 0.5 mL/minute flow rate. The resulting fractions were lyophilized to dryness and again monitored for the presence of the target band by HP-PAGE and alcian blue/silver staining. Fractions number 5 and 6 from the SEC were combined and re-separated one more time over the same Superdex Peptide column as described above. Lyophilized fractions number 6 and 7 from the second round of SEC (Figure S11B) were combined, applied to a Supelco Supelclean ENVI-Carb SPE graphite column (MilliporeSigma, Burlington, MA), and the target band eluted with 50% acetonitrile (Figure S11C) and lyophilized to dryness.

For NMR analysis, the graphite-column-purified, dried sample of the EPG-resistant target band was dissolved in 40 μL 75% D_2_O, 25% acetonitrile-d_3_ and transferred to a 1.7 mm NMR tube. Deuterated acetonitrile was added to this sample at 25% to help solvation of non-carbohydrate contaminants but it did not make a significant difference to most of the chemical shifts compared to the reference wine sample, which was in pure D_2_O (see below). ^13^C-HSQC data was collected at 25°C on an 800 MHz Bruker Neo spectrometer using the sequence hsqcetgpsisp2 from the standard Bruker library, with data sizes of 2048 x 256 points, acquisition times of 262 ms and 23 ms for t2 and t1, respectively, and 32 scans per t1 increment. The reference dried sample from wine was dissolved in 170 μL D_2_O, transferred to a 3 mm NMR tube, and data were collected on a 1100 MHz Bruker Neo spectrometer with similar acquisition parameters. Data were processed with Mnova software (Mestrelab, Inc).

### Direct Stochastic Optical Reconstruction Microscopy (d-STORM)

#### Sample preparation

The detailed immunohistochemistry protocol compatible with dSTORM imaging was previously published (Haas et al., 2020b; Haas et al., 2020a; Peaucelle et al., 2020). Samples were fixed in FAA buffer containing 50% ethanol, 10% acetic acid, and 5% formaldehyde for 1 hour at room temperature, then dehydrated sequentially in 70% ethanol, 95% ethanol, and 100% ethanol (twice), each for at least 30 minutes. Samples were cleared by incubating for 1 hour in 50% Histo-Clear and 50% ethanol, followed by 1 hour in 100% Histo-Clear. Next, samples were transferred to biopsy cassettes and infiltrated with paraffin: first in 50% Histo-Clear and 50% paraffin for 3 hours, then in 100% paraffin twice for 3 hours each, and finally overnight in 100% paraffin. Samples were stored at 4°C overnight before sectioning. Tissue was sectioned at a final thickness of 3-5 μm using a microtome and mounted on Ibidi microslides (Ibidi, catalog number: 80827). Deparaffinization was performed with three 30-minute washes in 100% Histo-Clear, followed by a 20-minute wash in 100% ethanol. Rehydration was performed through 15-minute incubations in 100% ethanol, 70% ethanol, 50% ethanol, 25% ethanol, 10% ethanol in 2F4 buffer, and finally 100% 2F4 buffer.

All remaining immunolabelling steps were performed using 2F4 buffer (20 mM Tris, 0.5 mM CaCl₂, 150 mM NaCl, pH 8). Free aldehyde groups were quenched using 50 mM NH₄Cl in 2F4 buffer for 15 minutes, followed by three washes with 2F4 buffer (each wash lasting 3–5 minutes). Primary antibodies were diluted in a blocking solution containing 1× 2F4 buffer and 5% dried milk. Primary and secondary antibodies were applied successively, each for 2 hours at room temperature or overnight at 4°C. Between each incubation step, samples were briefly washed three times in the blocking solution. After immunolabelling, samples were fixed in 3.7% formaldehyde in 1× 2F4 buffer for 10 minutes, washed briefly three times with 1× 2F4 buffer, then incubated for 15 minutes in 50 mM NH₄Cl in 2F4 buffer, followed by another three brief washes in 2F4 buffer. To label methylesterified HG, the LM20 rat monoclonal antibody was used at a 1:50 dilution. For calcium-coordinated demethylesterified HG, the 2F4 mouse monoclonal antibody was used at a 1:20 dilution. The following secondary antibodies were used: goat anti-mouse F(ab’)₂ fragment conjugated to Alexa Fluor 647 (Stratech Scientific, 115-607-003-JIR), diluted 1:1000; and goat anti-rat conjugated to CF568 (Sigma-Aldrich, SAB4600086), diluted 1:500.

#### Imaging

dSTORM imaging was performed in Glox buffer (Haas et al., 2020a) containing 50-100 mM MEA-HCl (Sigma, M6500), 10% glucose (Sigma), 0.5 mg/ml glucose oxidase (Sigma, G2133) and 40 μg/ml catalase (Sigma, C100) in 1×2F4 buffer at pH 7.5. Samples were imaged at room temperature in sealed 8-well Ibidi μ-slides on an inverted Nikon TiU2 Microscope equiped with Abbelight’s SAFE360 imaging system and operated by Abbelight NEO software. Samples were imaged in highly inclined illumination mode using Apochromat 100x/1.49 NA oil immersion objective. Typically, ∼35,000 frames were acquired using a Hamamatsu’s sCMOS Orca Flash BT camera with ∼50 ms frame acquisition time. Alexa647 dye was imaged with 640 nm, and CF568 with 561 nm excitation laser line. The 100 nm multicolor fluorescent microbeads were used as fiducial markers to register the two channels.

#### Data analysis

Single-molecule localization was performed using Abbelight NEO software. Lateral drift was automatically corrected using a correlation-based algorithm within the NEO software. For further analysis, molecular localization lists generated by NEO were imported into a custom-written, MATLAB-based single-molecule data analysis software available at https://github.com/inatamara/Grafeo-dSTORM-analysis- (Haas et al., 2020a; Peaucelle et al., 2020; Haas and Peaucelle, 2021).

Single-molecule data were filtered based on the number of photons (NP) emitted per molecule and the localization precision (LP). Further filtering was performed using Voronoi diagram (VD) thresholding by suppressing Voronoi Polygons (VP) larger than a threshold value. For cluster analysis, to retain only well-localized molecules, typical filtering thresholds were LP < 10 nm, NP > 1000, and VP < 10⁻⁹–10⁻⁸ nm³.

The coordinates of localized emitters were assigned to isolated clusters using 3D Voronoi diagrams and two-dimensional Delaunay triangulation (DT) with graph-based thresholding, as described previously (Haas et al., 2020a; Peaucelle et al., 2020; Haas and Peaucelle, 2021). The DT was represented as bidirectional graphs with points (nodes) connected by edges. Points were assigned to discrete cluster, connected component, by removing DT edges longer than ∼50 nm and eliminating graph nodes with a degree (number of connections) smaller than 5. The cluster length was estimated by extension—the maximum shortest paths spanning the DT graph (Haas et al., 2018).

All threshold values were adjusted slightly depending on signal noise and point density. Due to signal discontinuity, segmented clusters do not represent entire pectin nanofilaments but rather their segments. Statistical significance was assessed in MATLAB using the Kruskal–Wallis test for multiple group comparisons. *P*-values were adjusted using the Bonferroni correction.

## Acknowledgements

We thank Jiri Vlach for helpful discussions on RG-II-related structures. This material is based upon work supported by The BioEnergy Science Center (BESC), a US Department of Energy Bioenergy Research Center supported by the Office of Biological and Environmental Research in the Department of Energy’s Office of Science; by the Center for Bioenergy Innovation supported by the U.S. Department of Energy, Office of Science, Biological and Environmental Research under Contract Number ERKP886; and partially by the U.S. Department of Energy, Office of Science, Basic Energy Sciences, grant number DE-SC0015662 to the Complex Carbohydrate Research Center. This work was also supported by the European Research Council STORMtheWALL project 101041597 to K.T.H.

## Author Contributions

M.A.A. designed and performed most of the research, analyzed data, and wrote the paper. R.A.A. designed and performed HP-PAGE and chromatography, and analyzed data. K.T.H. and A.P. acquired and analyzed dSTORM data, prepared associated figures, and wrote the paper. J.G. and L.T. performed NMR experiments and analyzed data. I.M.B. and P.A. performed glycosyl linkage analysis and analyzed data. P.J.G., S.P. and S.K. carried out glycome profiling analysis. D.C.A., I.E., and J.G.I. performed research under supervision of M.A.A. S.E. helped in scanning electron microscopy sample preparation and acquisition. M.G.H. provided expertise and analyzed glycome profiling data. D.M. directed the research, analyzed the data and wrote the paper.

## Supplementary Data

**Supplementary Figure S1.** Heterozygous *GAUT1/gaut1-1* and complemented homozygous *gaut1-1/gaut1-1* plants grow normally in comparison to WT.

**Supplementary Figure S2.** Transcript expression of *GAUT1* and other *GAUT* genes in WT and *gaut1-1/gaut1-1* seedlings, as measured by quantitative RT-PCR.

**Supplementary Figure S3.** Arabidopsis eFP Browser (https://bar.utoronto.ca/efp_arabidopsis/cgi-bin/efpWeb.cgi) root high-resolution spatiotemporal transcript microarray data for *GAUT1* (Brady et al., 2007; Winter et al., 2007) showing highest levels of expression at the elongation zone.

**Supplementary Figure S4.** Arabidopsis WT, *GAUT1/gaut1-1*, and *gaut1-1/gaut1-1* calli.

**Supplementary Figure S5.** Growth of Arabidopsis WT, *GAUT1/gaut1-1*, and *gaut1-1/gaut1-1* suspension cells over 14 days of culture, as measured by (A) packed cell volume and (B) fresh weight.

**Supplementary Figure S6.** Glycosyl residue composition of extracellular material (ECM) secreted into the culture medium of 7-day old WT, *GAUT1/gaut1-1*, and *gaut1-1/gaut1-1* suspension cell cultures, as determined by GC-MS of trimethylsilyl (TMS) derivatives.

**Supplementary Figure S7.** Transcript expression of *GAUT1* and other *GAUT* genes in WT, *GAUT1/gaut1-1*, and *gaut1-1/gaut1-1* suspension culture cells over 14 days of culture, as measured by quantitative RT-PCR.

**Supplementary Figure S8.** Non-cellulosic glycosyl residue composition of total cell wall (destarched AIR), presented in mol%, of WT, *GAUT1/gaut1-1*, and *gaut1-1/gaut1-1* suspension cell culture lines, as determined by GC-MS of trimethylsilyl (TMS) derivatives.

**Supplementary Figure S9**. Analysis of wall fractions sequentially extracted from destarched AIR of WT, *GAUT1/gaut1-1* and *gaut1-1/gaut1-1* suspension culture lines using increasingly harsh solvents.

**Supplementary Figure S10.** High percentage polyacrylamide gel electrophoresis (HP-PAGE) comparison of the EPG-treated wall fractions from Arabidopsis WT suspension culture cells and RG-II monomer and dimer from wine.

**Supplementary Figure S11.** Chromatographic purification of the polymer band from WT EPG-digested 4M KOH fraction and visualization by HP-PAGE following alcian blue-silver nitrate-staining.

**Supplementary Figure S12.** Comparison of ^13^C-HSQC spectra of RG-II between the Arabidopsis sample isolated in this study, and an authentic sample from wine.

**Supplementary Figure S13.** HG:GalAT activity of microsomes extracted from WT, *GAUT1/gaut1-1*, and *gaut1-1/gaut1-1* suspension cultured cells harvested at different time points after the inception of the callus culture.

**Supplementary Table S1.** Arabidopsis *GAUT1* (*At3g61130*) T-DNA insertion mutant germplasm lines listed at The Arabidopsis Information Resource (TAIR) website and available from the Arabidopsis Biological Resource Center (ABRC).

**Supplementary Table S2.** PCR and quantitative RT-PCR primers used in this project

---

**Table S1.** Arabidopsis *GAUT1* (*At3g61130*) T-DNA insertion mutant germplasm lines listed at The Arabidopsis Information Resource (TAIR) website and available from the Arabidopsis Biological Resource Center (ABRC).

| Line | Stock name | *AtGAUT1*-associated polymorphism | T-DNA insertion location within *AtGAUT1* gene | T-DNA insertion | Remark |
| --- | --- | --- | --- | --- | --- |
| CS445115 | CS445115 | GK-470H07-019028 | Exon 10<sup>b</sup> | Double | Back-crossed to WT and screened over 4 generations to isolate line with single T-DNA insertion within the *GAUT1* gene that is used for analysis in this study and re-named *gaut1-1*. |
| SALK_130652 | SALK_130652 | SALKSEQ_130652.1 | Intron 2<sup>a,b</sup> | Triple | Identified stunted *gaut1*<sup>-/-</sup> seedlings similar to those of CS445115; renamed *gaut1-3*. |
| SALK_063880 | SALK_063880 | SALKSEQ_063880.1 | Exon 10<sup>a</sup> | Triple | Identified stunted *gaut1*<sup>-/-</sup> seedlings similar to those of CS445115; renamed *gaut1-2*. |
| SAIL_1162_A03 | CS878267 | SAIL_1162_A03 | 5'-UTR<sup>a</sup> | Single | Only WT was identified by genotyping (26/26)<sup>c</sup> |
| WiscDsLoxHs190_06B | CS918178 | WISCDSLOXHS190_06B.1 | 5'-UTR<sup>b</sup> | Single | Isolated HM looks comparable to WT. |

<sup>a</sup> T-DNA insertion location was determined based on information available at the TAIR website on the insertion flanking sequence that spans the insertion junction and thus encompasses both the *GAUT1* gene and the T-DNA sequences.
<sup>b</sup> T-DNA insertion location was determined based on sequencing of product from PCR reaction using the gene specific and the T-DNA primers that amplify the region spanning the insertion junction.
<sup>c</sup> Values in parentheses indicate the numbers of WT plants over the numbers of plants genotyped.

**Table S2.**
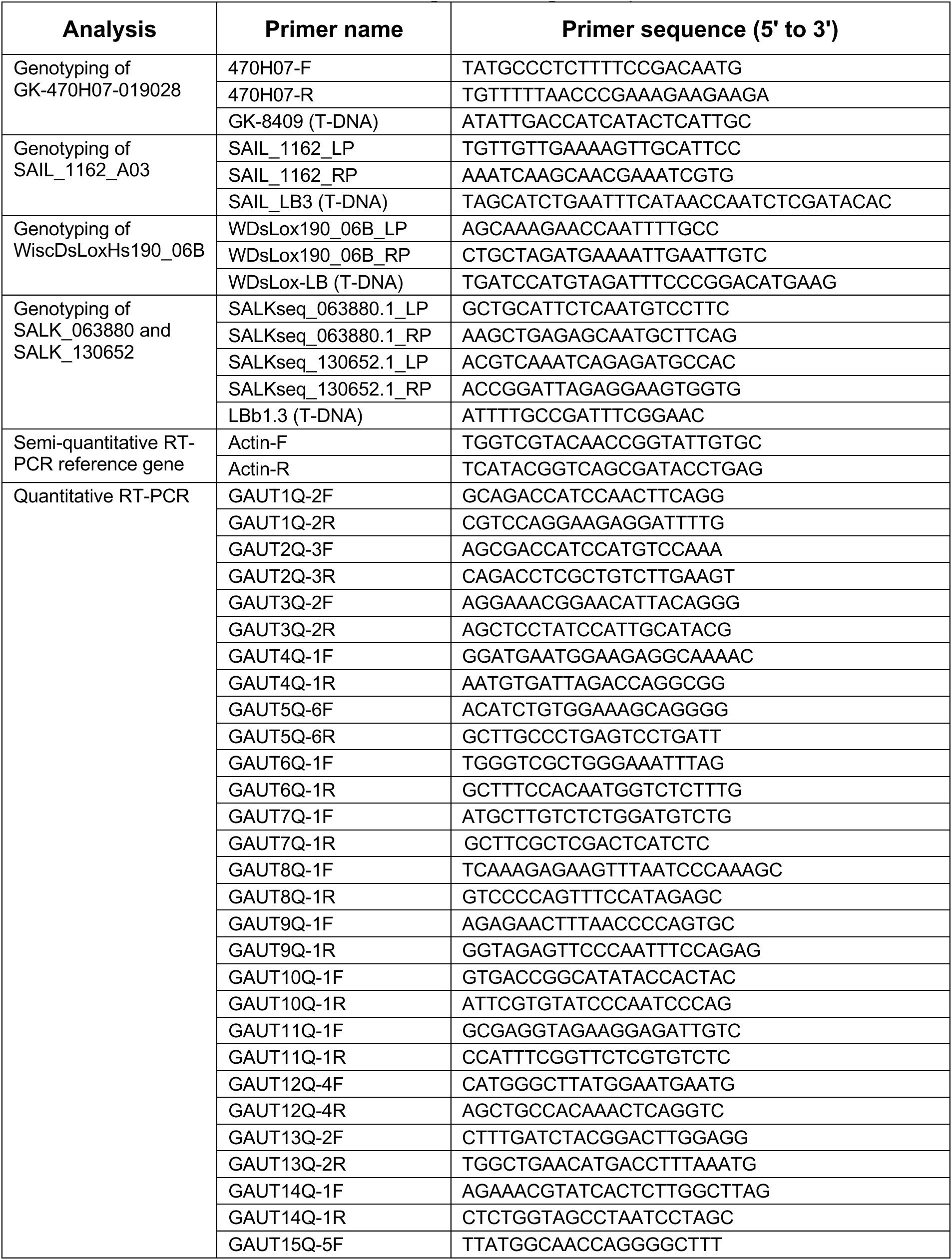
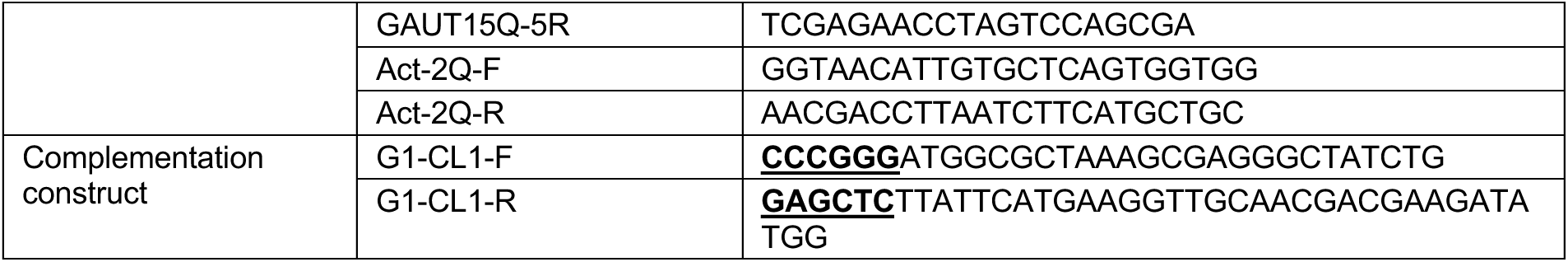
PCR and quantitative RT-PCR primers used in this project. For complementation construct primers, in bold and underlined are the added restriction sites for SmaI and SacI for the forward and reverse primers, respectively.

