## Supplementary Figures for "Arabidopsis GALACTURONOSYLTRANSFERASE (GAUT) 1 synthesizes a homogalacturonan tightly bound to the cell wall and required for cell expansion"

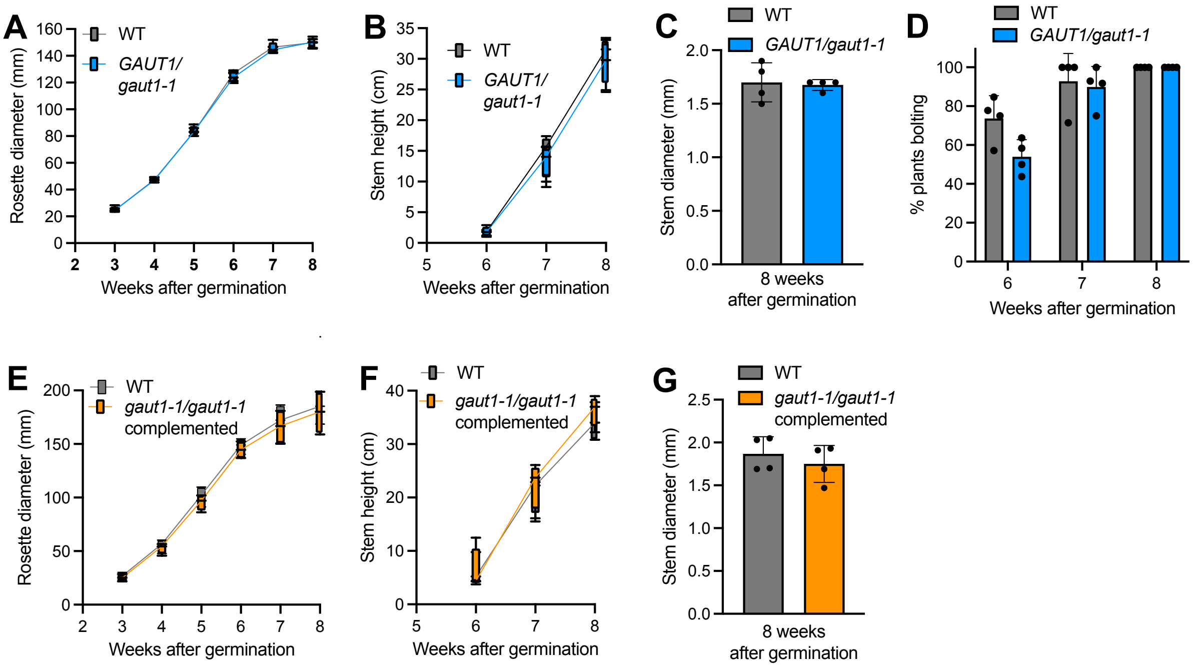


**Figure S1. Heterozygous *GAUT1/gaut1-1* and complemented homozygous *gaut1-1/gaut1-1* plants grow normally in comparison to WT.**

(A-D) Heterozygous *GAUT1/gaut1-1* plants have comparable (A) rosette diameter, (B) stem height, (C) stem diameter (measured at 8 weeks old), and (D) flowering time compared to WT. (E-G) Complemented homozygous *gaut1-1/gaut1-1* plants showed comparable (E) rosette diameter, (F) stem height, and (G) stem diameter (measured at 8 weeks old) compared to WT. Data are means ± standard deviation from four independent experiments (n=4) for a total of >40 plants for A-D, and >60 plants for E-G. No significant difference was observed as analyzed by Student's t-test ($P > 0.05$).

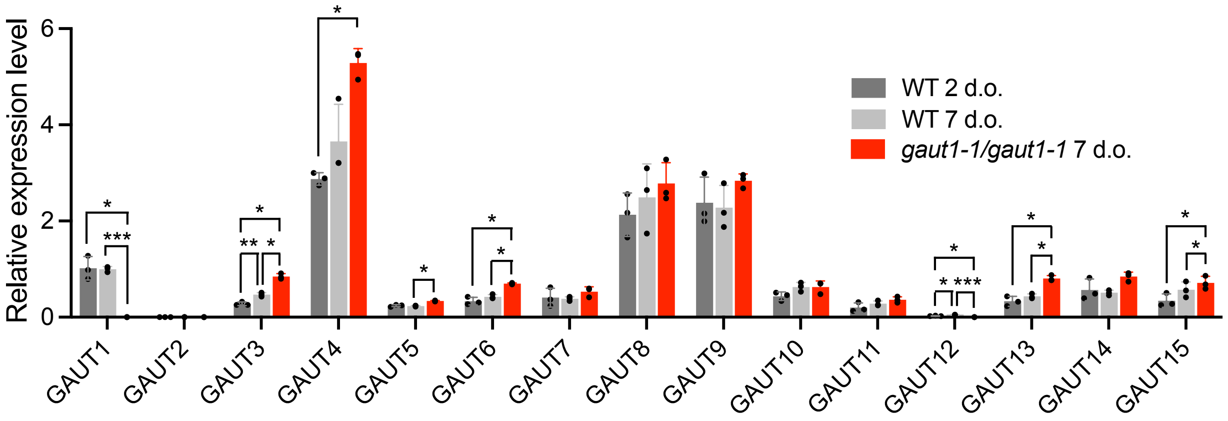


**Figure S2. Transcript expression of *GAUT1* and other *GAUT* genes in WT and *gaut1-1/gaut1-1* seedlings, as measured by quantitative RT-PCR.**

Expression of *GAUT1* in 2-day-old WT seedlings was set to 1. Data are means ± standard deviation from three independent experiments (n=3). Asterisks indicate significant difference as determined by ANOVA followed by Tukey's multiple comparison test (* $P < 0.05$, ** $P < 0.01$, *** $P < 0.001$).

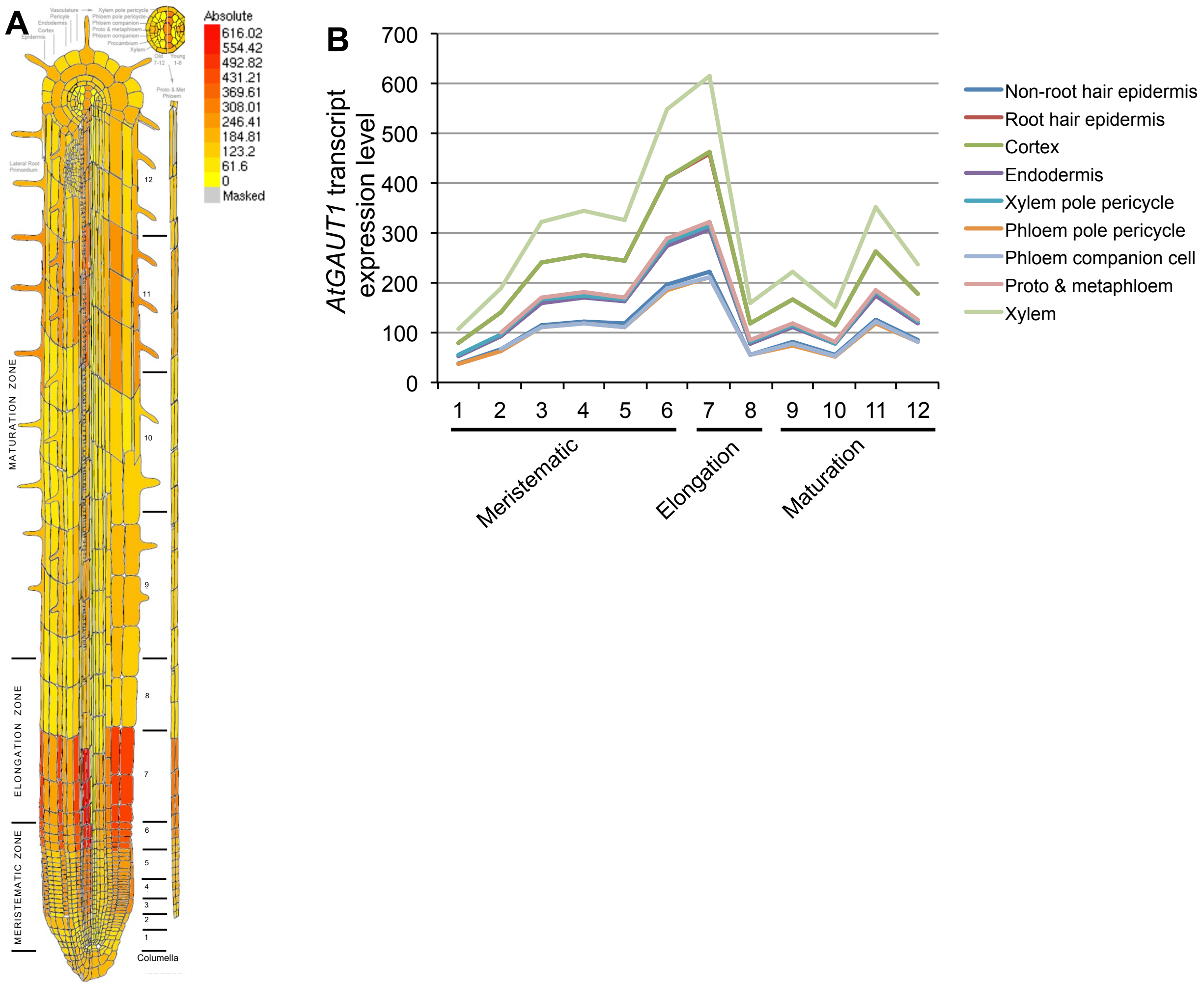


**Figure S3. Arabidopsis eFP Browser ([https://bar.utoronto.ca/efp_arabidopsis/cgi-bin/efpWeb.cgi](https://bar.utoronto.ca/efp_arabidopsis/cgi-bin/efpWeb.cgi)) root high-resolution spatiotemporal transcript microarray data for *GAUT1* (Brady et al., 2007; Winter et al., 2007) showing highest levels of expression at the elongation zone.**

(A) Graphical representation, and (B) plot of longitudinal expression raw data.

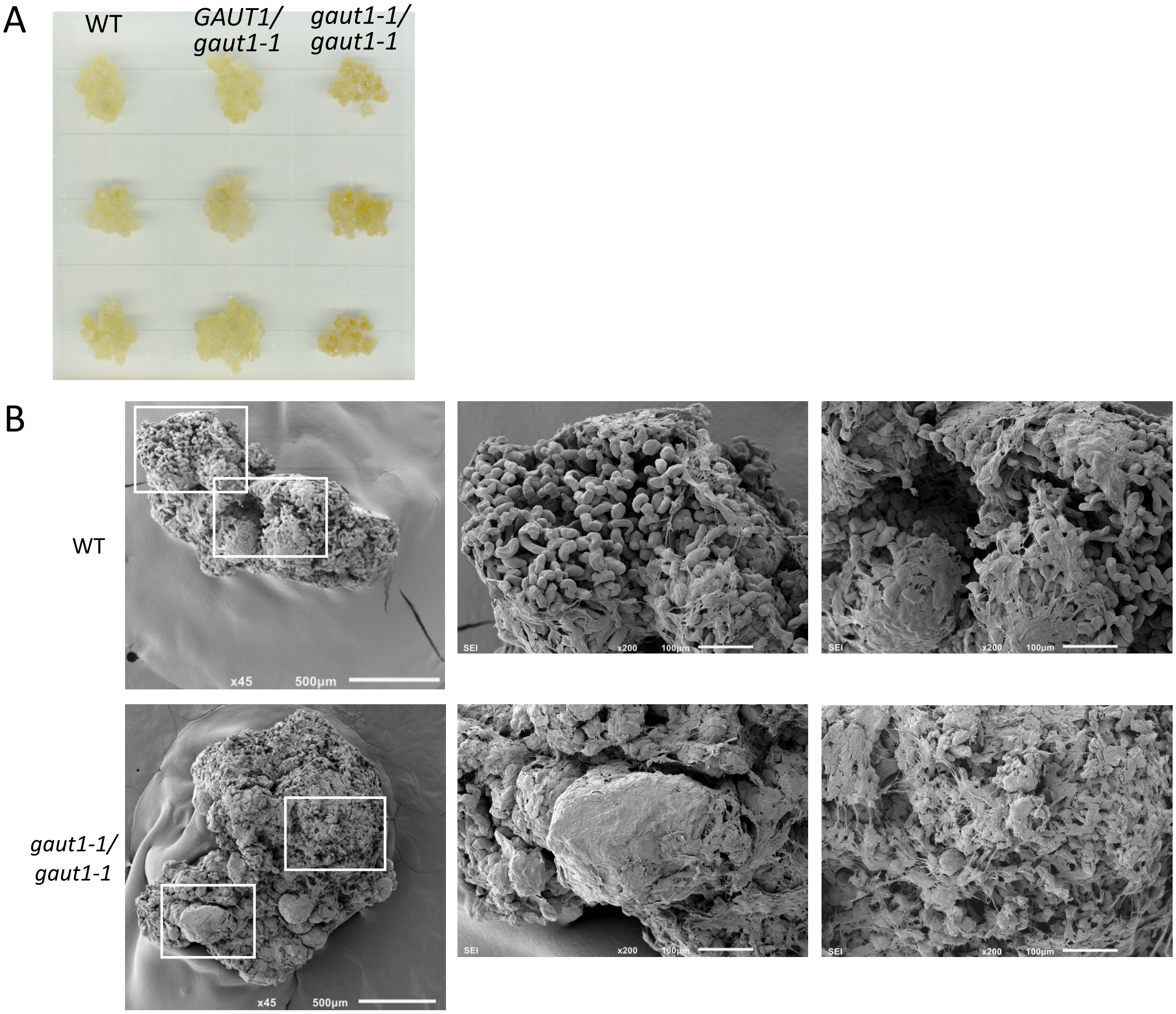


**Figure S4. Arabidopsis WT, *GAUT1/gaut1-1*, and *gaut1-1/gaut1-1* calli.**
(A) A representative photograph of WT, *GAUT1/gaut1-1*, and *gaut1-1/gaut1-1* calli on a medium plate. (B) Representative scanning electron micrographs of the calli, showing how the extracellular polymer covers the callus surface more extensively in the *gaut1-1/gaut1-1* line compared to WT. Areas delineated by the white boxes in the left-hand panels (scale bars: 500 µm) are shown at higher magnification in the middle and right-hand panels (scale bars: 100 µm).

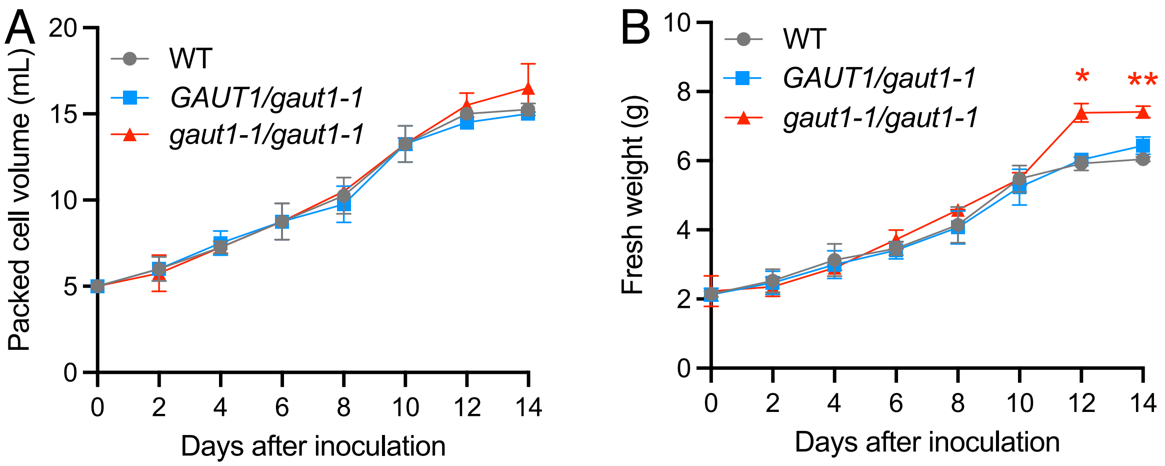


**Figure S5. Growth of Arabidopsis WT, *GAUT1/gaut1-1*, and *gaut1-1/gaut1-1* suspension cells over 14 days of culture, as measured by (A) packed cell volume and (B) fresh weight.** Five mL of packed 10-day-old suspension cells were used to inoculate 100 mL medium in 500 mL erlenmeyer flasks. Cultures were incubated with constant shaking at 100 rpm at room temperature in the dark. Data are means ± standard deviation from two independent experiments (n=2). Asterisks indicate significant difference to WT as analyzed by Analysis of Variance (ANOVA) followed by Tukey's multiple comparison test (* $P < 0.05$, ** $P < 0.01$).

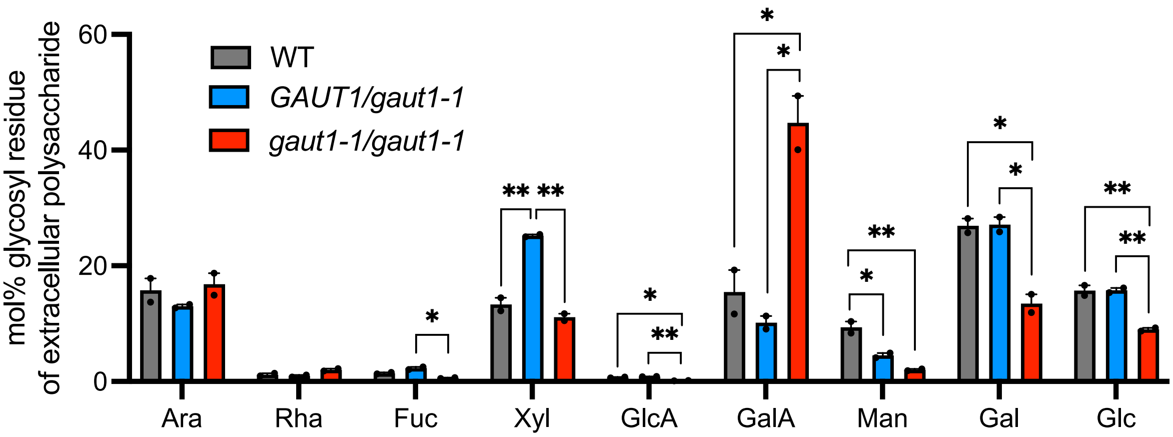


**Figure S6. Glycosyl residue composition of extracellular material (ECM) secreted into the culture medium of 7-day old WT, *GAUT1/gaut1-1*, and *gaut1-1/gaut1-1* suspension cell cultures, as determined by GC-MS of trimethylsilyl (TMS) derivatives.**
Data are means ± standard error of three technical replicates of ECM extracted from the medium of two independent culture batches (n=2). Asterisks indicate significant difference as analyzed by one-way ANOVA followed by Tukey's multiple comparison test (* $P < 0.05$, ** $P < 0.01$, *** $P < 0.001$).

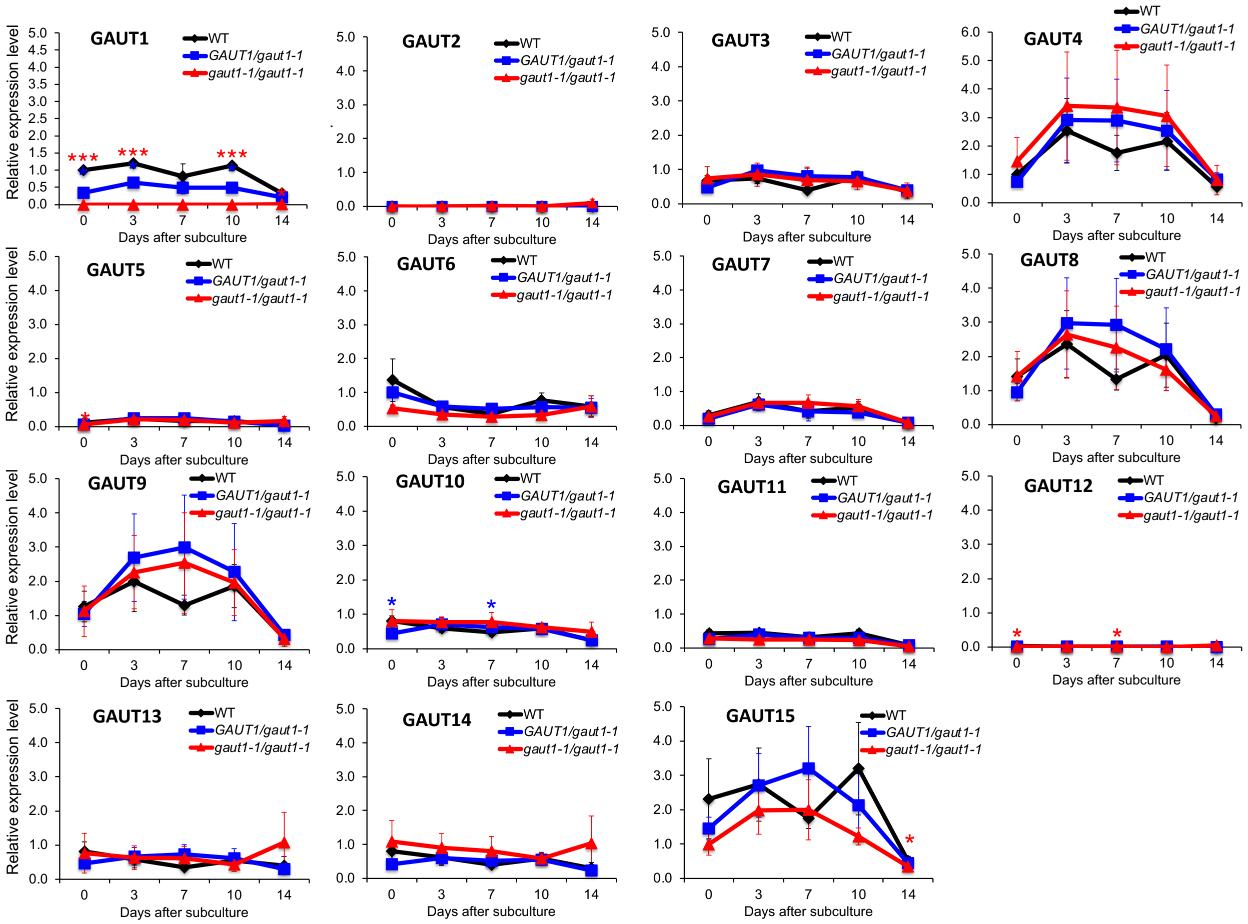


**Figure S7. Transcript expression of *GAUT1* and other *GAUT* genes in WT, *GAUT1/gaut1-1*, and *gaut1-1/gaut1-1* suspension culture cells over 14 days of culture, as measured by quantitative RT-PCR.**

Expression of *GAUT1* in WT cells at day 0 is set to 1. Data are means ± standard deviation from three independent experiments (n=3). Asterisks indicate significant difference compared to WT, as determined by ANOVA followed by Tukey's multiple comparison test (* $P < 0.05$, *** $P < 0.001$).

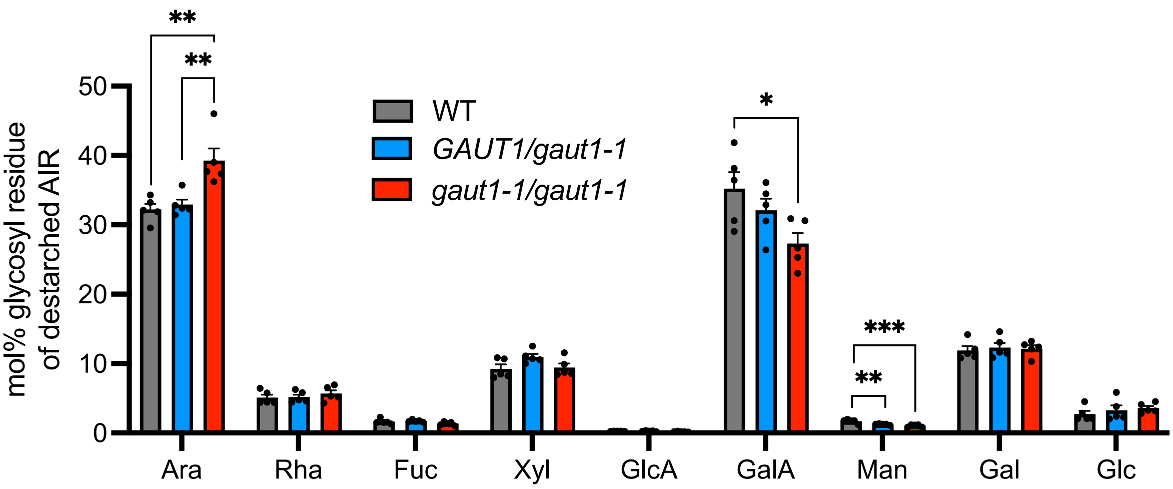


**Figure S8. Non-cellulosic glycosyl residue composition of total cell wall (destarched AIR), presented in mol%, of WT, *GAUT1/gaut1-1*, and *gaut1-1/gaut1-1* suspension cell culture lines, as determined by GC-MS of trimethylsilyl (TMS) derivatives.**

Data are means ± standard error of two technical replicates of destarched AIR extracted from five independent culture batches (n = 5). Asterisks indicate significant difference as analyzed by one-way ANOVA followed by Tukey's multiple comparison test (* $P < 0.05$, ** $P < 0.01$, *** $P < 0.001$).

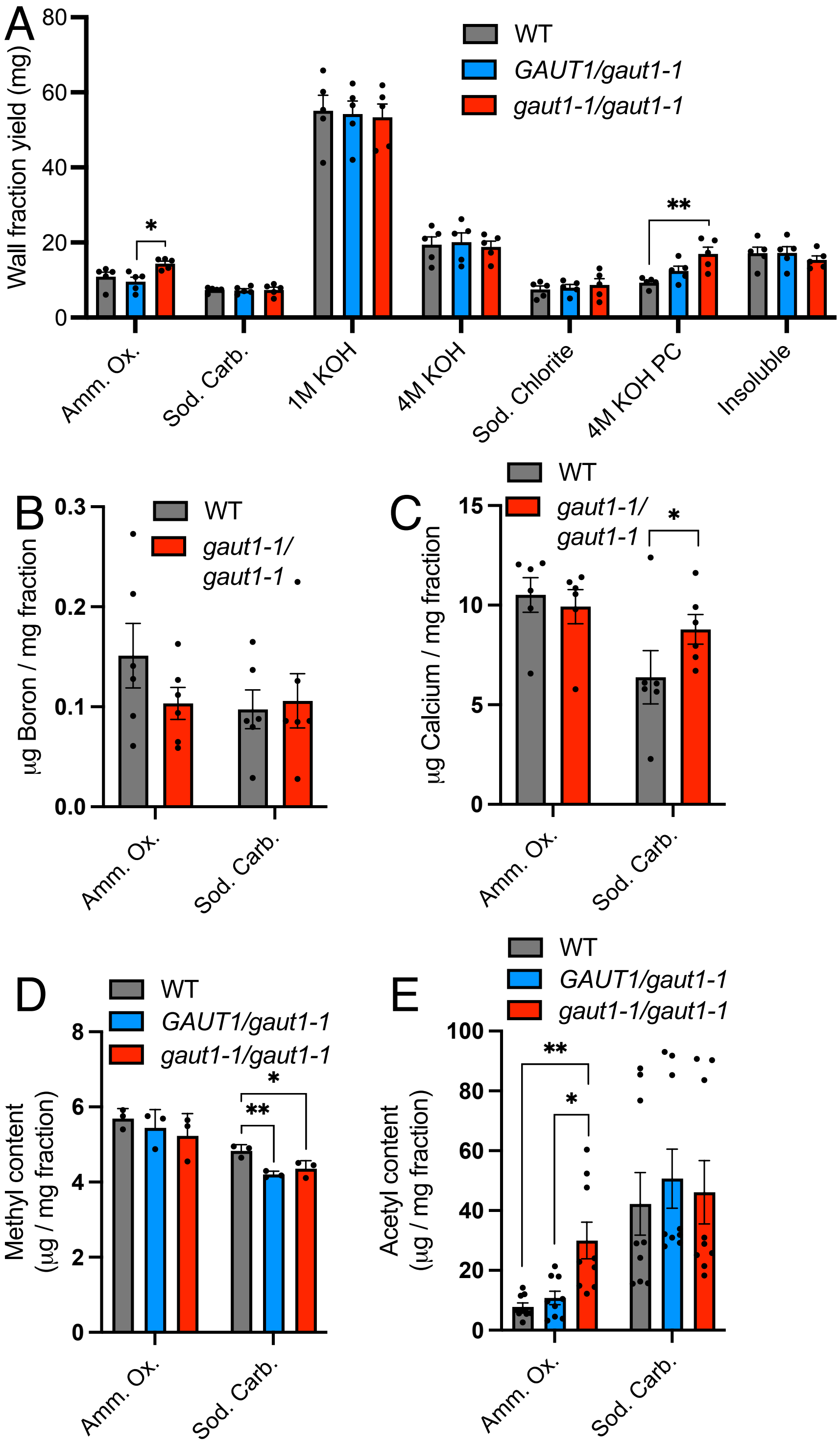


**Figure S9. Analysis of wall fractions sequentially extracted from destarched AIR of WT, *GAUT1/gaut1-1* and *gaut1-1/gaut1-1* suspension culture lines using increasingly harsh solvents.**
(A) Mass of each wall fraction sequentially extracted from 200 mg of destarched AIR using the following solvents: 50 mM ammonium oxalate (Amm. Ox.); 50 mM sodium carbonate (Sod.

Carb.); 1M KOH; 4M KOH; 100 mM sodium chlorite (Sod. Chlorite); 4M KOH post-chlorite (4M KOH PC). Data are means ± standard error from duplicate extractions of three independent culture batches (n=6). (B) Boron, (C) calcium, (D) methyl, and (E) acetyl content of ammonium oxalate and sodium carbonate fractions. Data are means ± standard error from extracts of three independent culture batches (n=3). Asterisks indicate statistically significant difference as determined by ANOVA followed by Tukey's multiple comparison test (* $P < 0.05$, ** $P < 0.01$, *** $P < 0.001$).

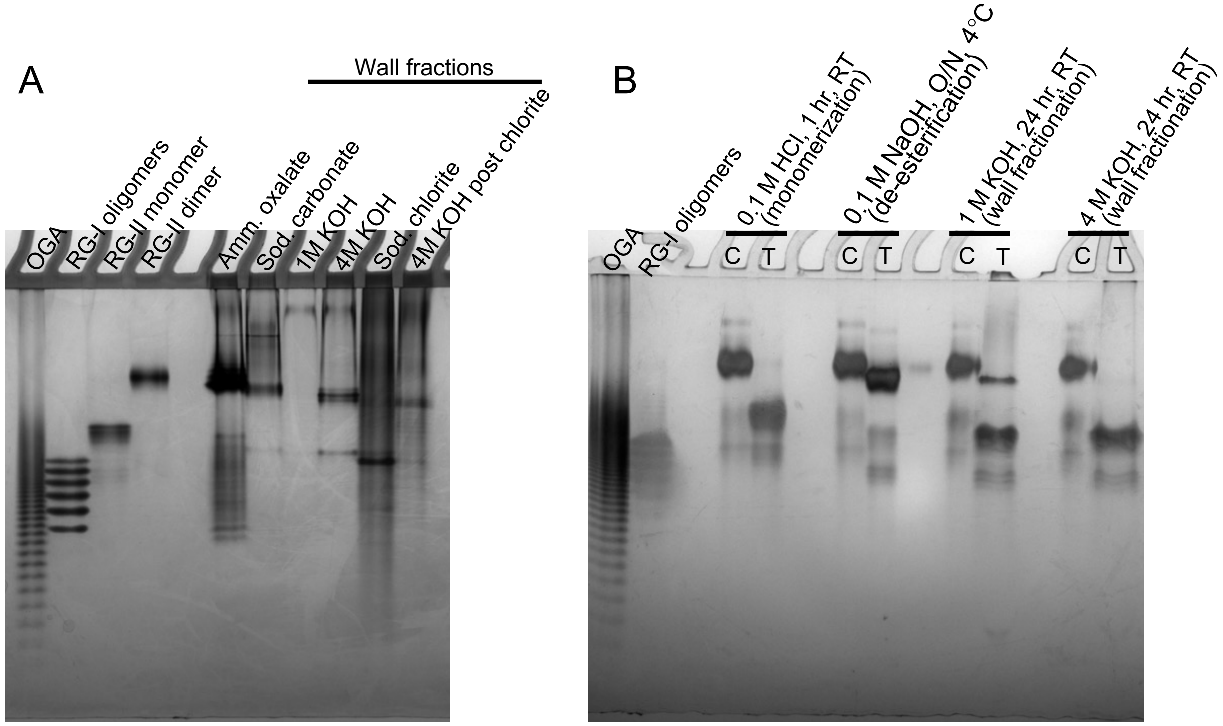


**Figure S10. High percentage polyacrylamide gel electrophoresis (HP-PAGE) comparison of the EPG-treated wall fractions from Arabidopsis WT suspension culture cells and RG-II monomer and dimer from wine.**

(A) Alignment of standard RG-II monomer and dimer from wine with the EPG-treated wall fractions from Arabidopsis WT suspension cultured cells. Note that Arabidopsis RG-II was reported to lack the terminal β-L-Ara*f* and the α-L-Rha*p* attached to the O-3 of Ara*p* of side chain B, thus overall it is smaller in size compared to RG-II from wine (Glushka et al., 2003). It can be observed from the gel picture that (1) the bulk of RG-II is extracted as dimers in the ammonium oxalate fraction and to a lesser extent in virtually all fractions as dimers and/or monomers; (2) Sodium carbonate and subsequent extraction steps appears to gradually de-esterify and/or degrade the RG-II dimer and monomer. OGA – oligogalacturonides.

(B) The effect of different treatments on RG-II dimer standard from wine (C – Control, T – Treatment): RG-II monomerization by 0.1 M HCl for 1 hr at room temperature (RT); RG-II de-esterification by 0.1 M NaOH for overnight (O/N) at 4°C; the third step of wall sequential extractions by 1 M KOH (with 1% sodium borohydride) for 24 hr at RT; and the fourth step of wall sequential extractions by 4 M KOH (with 1% sodium borohydride) for 24 hr at RT. Note that de-esterification with 0.1 M NaOH shifted the migration of the RG-II dimer band on the HP-PAGE due to change in charge and/or size. Note as well that the 1 M KOH treatment appears to de-esterify and partially monomerize the RG-II dimer, while the 4 M KOH treatment completely de-esterify and monomerize the RG-II dimer.

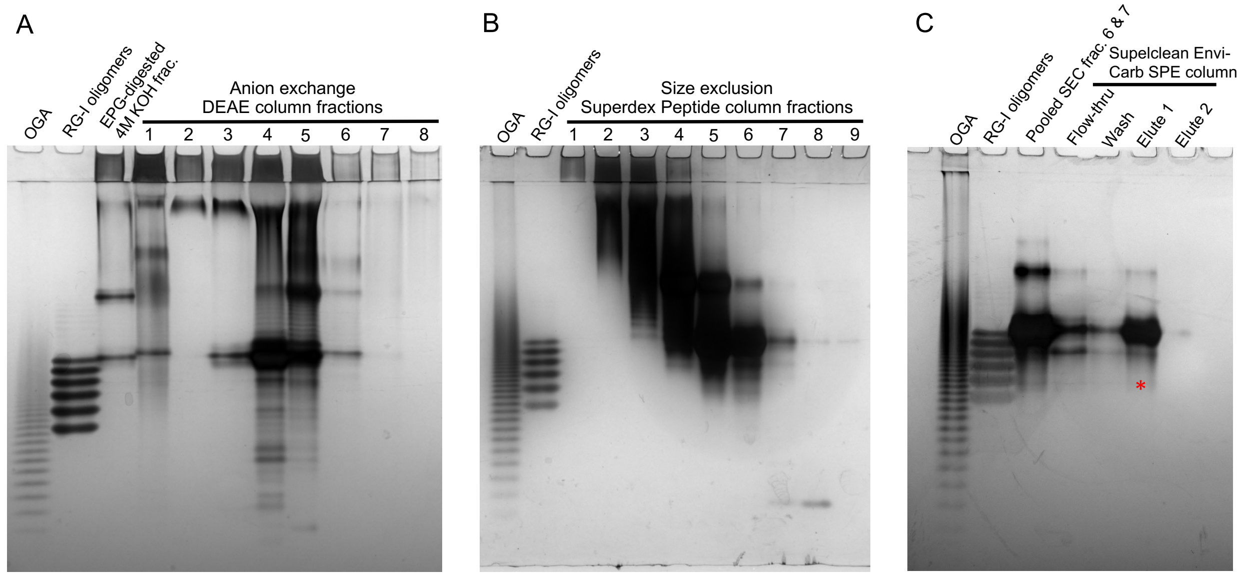


**Figure S11. Chromatographic purification of the polymer band from WT EPG-digested 4M KOH fraction and visualization by HP-PAGE following alcian blue-silver nitrate-staining.**

(A) Anion exchange chromatography over a DEAE Sephacel column with elution by an ammonium formate (AF) step gradient as follows: fraction 1, 50 mM AF; fraction 2, 100 mM AF; fraction 3, 200 mM AF; fraction 4, 300 mM AF; fraction 5, 400 mM AF; fraction 6, 500 mM AF; fraction 7, 2 M AF. Fraction 8 was eluted with 2 M NaCl. Fractions 3-6 containing the polymer band of interest were pooled.

(B) Size exclusion chromatography of the pooled DEAE fractions 3-6 over a Superdex Peptide 10/300 GL column with isocratic elution in 50 mM ammonium formate for 60 minutes, flow rate 0.5 mL/minute. Fractions were collected as follows: fraction 1, 12-16 minutes; fraction 2, 16-18 minutes; fraction 3, 18-20 minutes; fraction 4, 20-22 minutes; fraction 5, 22-24 minutes; fraction 6, 24-26 minutes; fraction 7, 26-28 minutes; fraction 8, 28-30 minutes; and fraction 9, 30-35 minutes. Fractions 6 and 7 were pooled.

(C) Solid phase extraction of the pooled SEC fractions 6 and 7 over a Supelco Supelclean ENVI-Carb 3-mL graphite column. Eluents used (2 mL each): Wash – ddH2O, Elute 1 – 50% acetonitrile, Elute 2 – 100% acetonitrile. Red asterisk denotes the Elute 1 fraction that was subsequently subjected to NMR analysis.

Standards: OGA, mixture of oligogalacturonides enriched for DP 7-23; RG-I oligomers, mixture of RG-I backbone oligomers DP 12-22.

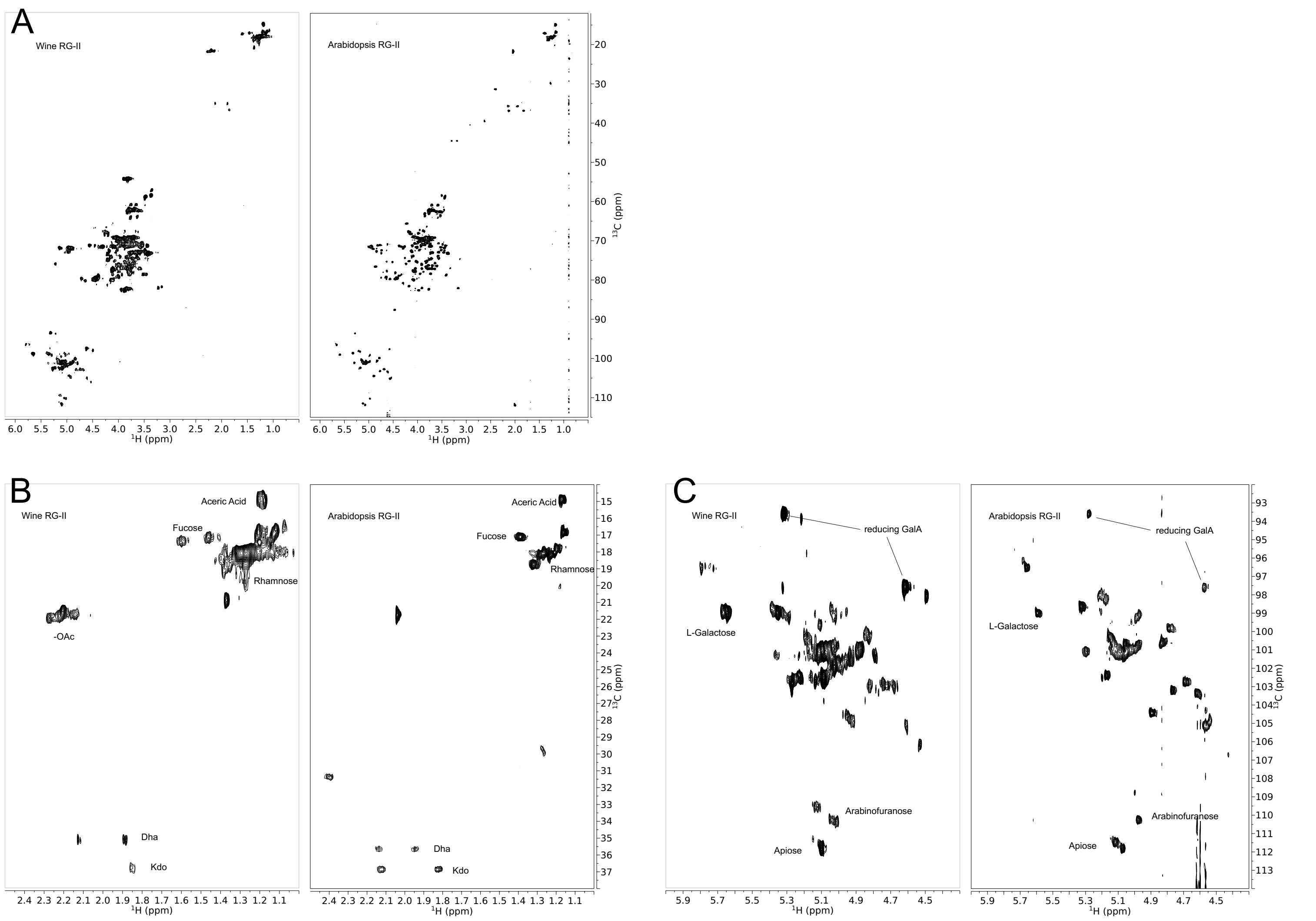


**Figure S12. Comparison of $^{13}$C-HSQC spectra of RG-II between the Arabidopsis sample isolated in this study, and an authentic sample from wine.** (A) Full carbohydrate region, (B) methyl and methylene region, and (C) anomeric region. Some signals are identified by residue type based on Herve du Penhoat et al. (1999), Glushka et al. (2003), and Rodriguez-Carvajal et al. (2003).

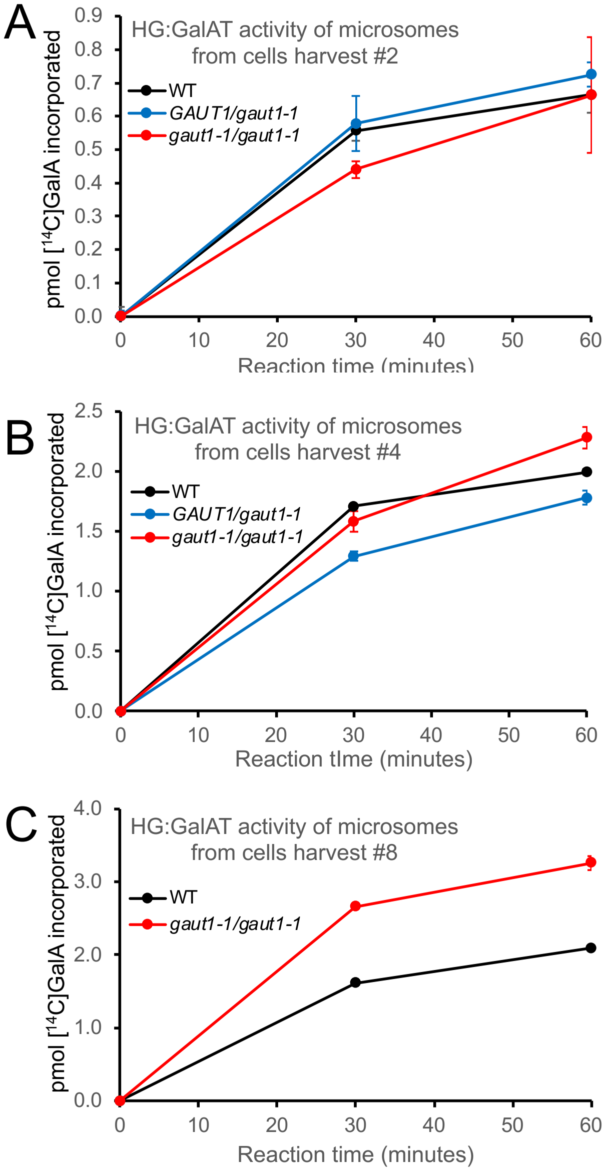


**Figure S13. HG:GalAT activity of microsomes extracted from WT, *GAUT1/gaut1-1*, and *gaut1-1/gaut1-1* suspension cultured cells harvested at different time points after the inception of the callus culture.** (A) Cell harvest #2, 11 months old; (B) cell harvest #4, 1 year and 6 months old; (C) cell harvest #8, 2 years and 4 months old.
